# Establishment of a new sex-determining allele driven by sexually antagonistic selection

**DOI:** 10.1101/2020.10.01.322412

**Authors:** T. Sakamoto, H. Innan

**Affiliations:** SOKENDAI, The Graduate University for Advanced Studies, Hayama, Kanagawa 240-0193, Japan

**Keywords:** diffusion theory, sex locus, population genetics, establishment

## Abstract

Some species undergo frequent turnovers of sex-determining locus, rather than having stable diverged sex chromosomes. In such species, how often turnover occurs is a fundamental evolutionary question. We model the process with considering a linked locus under sexually antagonistic selection. The entire process of a turnover may be divided into two phases, which are referred to as the stochastic and deterministic phases. The stochastic phase is when a new sex-determining allele just arises and is still rare and random genetic drift plays an important role. In the deterministic phase, the new allele further increases in frequency by positive selection. The theoretical results currently available are for the deterministic phase, which demonstrated that a turnover of a newly arisen sex determining locus could benefit from selection at a linked locus under sexually antagonistic selection, by assuming that sexually antagonistic selection works in a form of balancing selection. In this work, we provide a comprehensive theoretical description of the entire process from the stochastic phase to the deterministic phase. In addition to balancing selection, we explore several other modes of selection on the linked locus. Our theory allows us make a quantitative argument on the rate of turnover and the effect of the mode of selection at the linked locus. We also performed simulations to explore the pattern of polymorphism around the new sex determining locus. We find that the pattern of polymorphism is informative to infer how selection worked through the turnover process.

---

**R**ecent genome analyses have demonstrated that genetic systems that determine sex are more labile than previously thought and that the turnover of sex-determining loci has repeatedly occurred. In some clades such as teleost fish and amphibians, sex-determining loci differ among closely related species or even within a species (Bachtrog *et al*. 2014; Beukeboom and Perrin 2014). Frequent turnover should occur in species especially when sex is determined by a single sex-determining locus, rather than a pair of highly diverged sex chromosomes. Turnover should be initiated by mutation at another potentially sex-determining locus, which could become a new sex-determining locus while the dimorphism at the original sex-determining locus disappears.

Many theoretical studies have investigated the evolutionary process of such turnover (Bull and Charnov 1977; van Doorn and Kirkpatrick 2007, 2010; Kozielska *et al*. 2010; Blaser *et al*. 2013, 2014; Veller *et al*. 2017; Scott *et al*. 2018; Saunders *et al*. 2018, 2019). A consensus has been established that, if a new sex-determination system has higher fitness than the old one, the new system could potentially override the old one. This explains why turnover hardly occurs in species with a diverged pair of sex chromosomes, such as the X/Y system in mammals and the W/Z system in birds. Theoretical examinations of turnover usually involves a two-locus system, under which turnover has to pass through a phase in which dimorphic sex-determining alleles segregate at both of the two sex-determining loci. A deterministic theory assuming a population with an infinite size (Bull and Charnov 1977) demonstrated that the system with higher fitness can be stably maintained by selection. It is indicated that the fitness advantage of the new locus over the existing one would be an important factor for the turnover of sex-determining loci.

A possible scenario that a new sex-determining locus confers a fitness advantage is that the new locus arises in close linkage to a locus under sexually antagonistic selection (van Doorn and Kirkpatrick 2007, 2010). This process is explained in Figure 1A. Initially, sex is determined by the original X/Y locus, while the A/a locus is another potential sex-determining locus on a different chromosome (autosome). That is, the A/a locus is monomorphic (fixed for allele a) so that it does not play a role in sex determination, but when allele A arises from allele a by mutation, it creates a new sex-determination system. We refer to allele A as a sex-determining allele. There is another polymorphic locus (B/b) that is located close to the A/a locus. The B/b locus is assumed to be subject to sexually antagonistic selection; for example, allele B is beneficial in males and allele b is beneficial in females. At this point, the B/b locus does not confer any advantage or disadvantage because there is no physical linkage to the sex-determining locus X/Y. If a sex-determining allele (A) arises by mutation at the A/a locus, then one of the possible outcomes is that this new dimorphic locus takes over the role of sex determination and the original X/Y locus becomes monomorphic.

**Figure 1.**
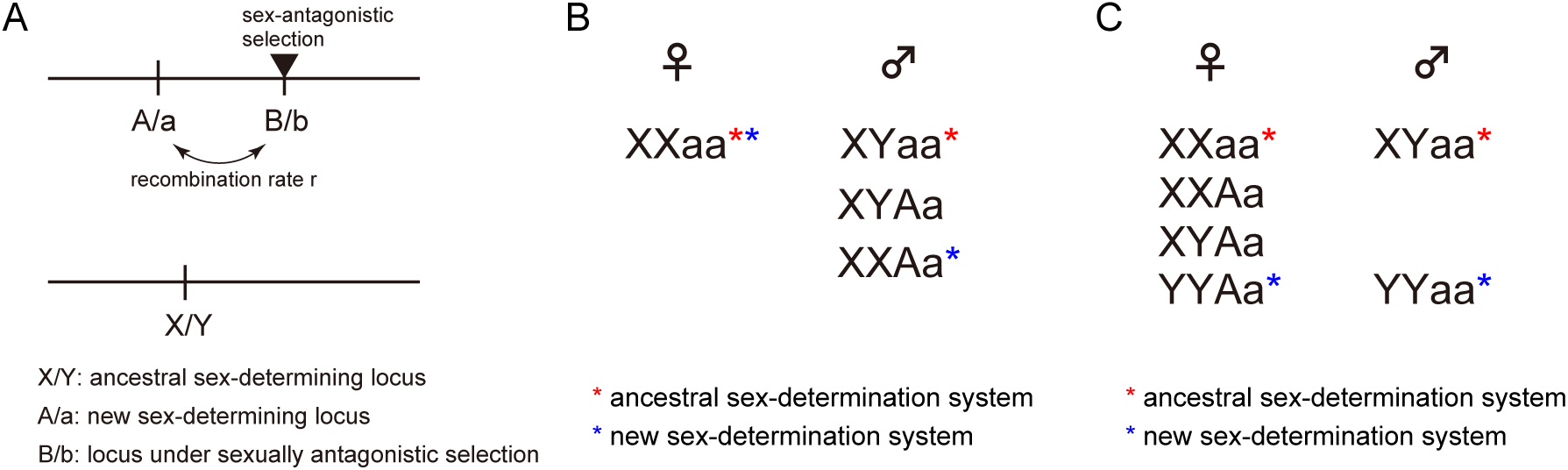
(A) The three loci model used in this work. (B) Relationship between sexes and genotypes when allele A has a masculinizing effect. (C) Relationship between sexes and genotypes when allele A has a feminizing effect. The red and blue stars are given to the genotypes that determine sex under the ancestral and new systems, respectively. The genotypes with no stars arise in the phase of transition from the ancestral to the new system.

We are here interested in how often such turnover of sex-determining loci occurs. To understand this, it is crucial to theoretically describe the entire process, from the birth of a new sex-determining allele to its stable establishment. However, previous studies on this topic mainly by van Doorn and Kirkpatrick (2007, 2010) focused on the second half of the process (as explained below). The purpose of this work is to provide an analytical description for the first half, which largely determines how often turnover occurs under what conditions.

The entire process may be divided into two phases, which are referred to as the stochastic and deterministic phases. The stochastic phase starts when a new sex-determining allele A arises, and continues until the frequency becomes high enough to escape from extinction due to random genetic drift. Then, the deterministic phase follows, in which allele A further increases in frequency by positive selection and genotype Aa becomes fixed in the heterogametic sex; in this way, the new A/a locus takes over the role in sex determination. The theoretical results currently available are for the deterministic phase, in which analytical treatment is quite straightforward because random genetic drift may be ignored. van Doorn and Kirkpatrick (2007, 2010) used a deterministic approach to describe the rate of increase in the frequency of allele A during the early stages of the deterministic phase. In contrast, the behavior of allele A in the initial stochastic phase is much more complicated and many factors are involved in it, which is the scope of this work. One such factor is the effect of random genetic drift, which is the major evolutionary force acting when the allele frequency is low. The linkage to the B/b is also a very important factor. If allele A arises in linkage with allele B, allele A immediately benefits from positive selection because the haplotype A-B confers a selective advantage. On the other hand, if allele A arises in linkage with allele b, selection works against allele A because the haplotype A-b is deleterious. Recombination plays a crucial role in determining the rate at which the advantageous haplotype A-B is created and broken up. Another important factor is how selection works on the B/b locus. We here provide a comprehensive theoretical description of this complex process in the stochastic phase, which contributes to a full understanding of the microevolutionary process of the turnover of sex-determining loci together with the theory for the deterministic phase.

Through this work, we find that the mode and intensity of selection at the B/b locus are particularly important in the stochastic phase, in which the fate of the new mutation is determined. Depending on the parameter setting for the effect of the B/b locus on fitness, selection operates in various modes (see Figure 2): Selection works for or against allele B, or even in some parameter space, balancing selection works to maintain the two alleles at intermediate frequencies. It is easy to imagine that this parameter setting largely affects the fate of the sex-determining mutation. However, perhaps because the stochastic phase was not the major focus, van Doorn and Kirkpatrick (2007, 2010) mainly focused on the parameter space where the B/b alleles are maintained in intermediate frequencies by balancing selection (i.e., Figure 2F), and demonstrated that turnover of the A/a locus likely occurs when there is tight linkage to the B/b locus. Note that balancing selection can operate only in a relatively narrow parameter space of the fitness effect of the B/b locus (see below). Furthermore, there are no strong biological reasons why balancing selection works on sexually antagonistic alleles (van Doorn and Kirkpatrick 2007, 2010). In this work, we explore more general situations with various modes of selection, which results in a quantitatively different conclusion from that in the previous work (van Doorn and Kirkpatrick 2007, 2010).

**Figure 2.**
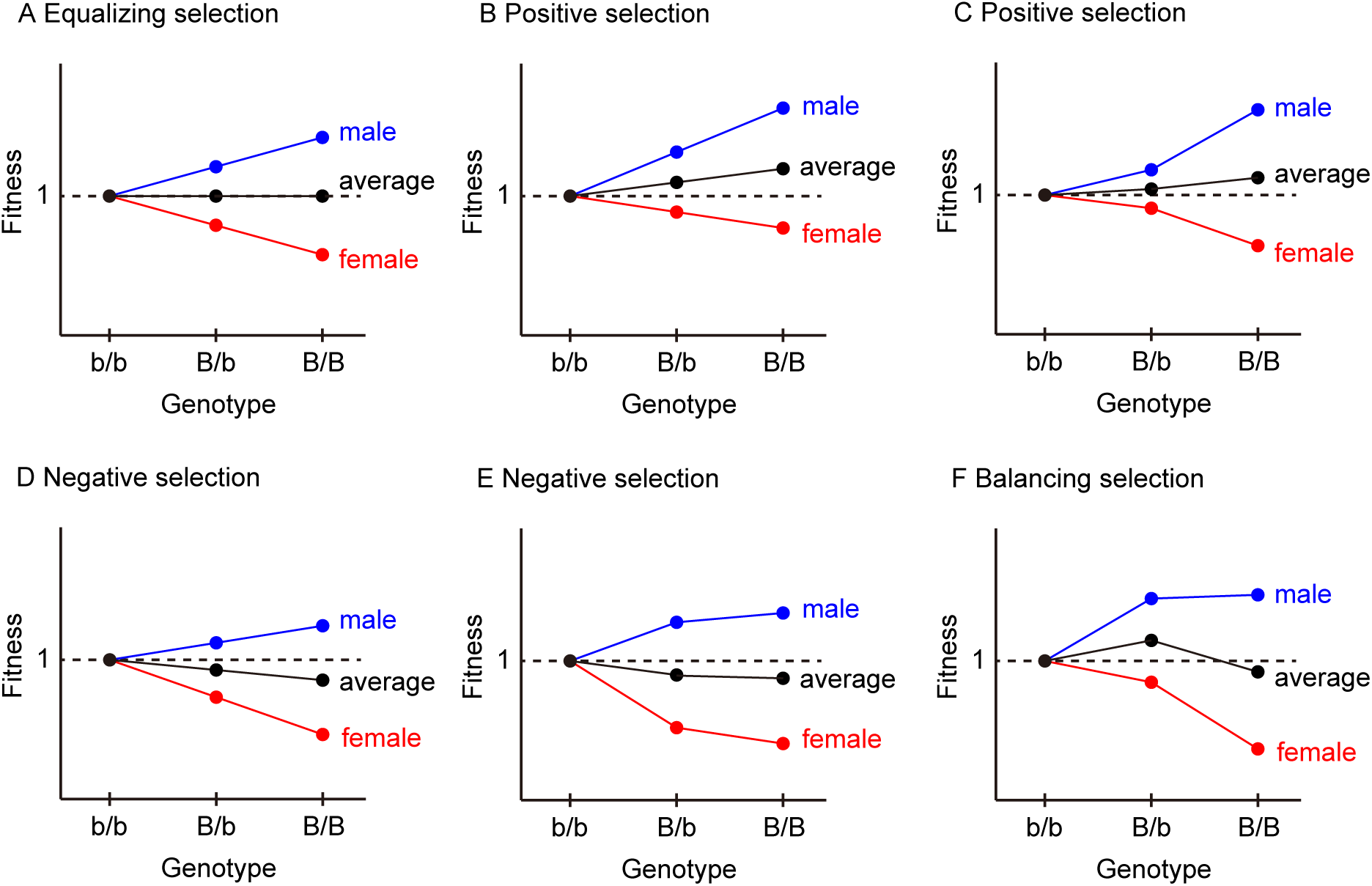
Various modes of selection depending on dominance and selection coefficient at the B/b locus when locus X/Y determines sex. The net effect of selection at locus B/b is given by the average of the two sexes. If the dominance and strength of selection are similar in males and females but the directions of selection are opposite, the fitness of allele B equals the fitness of allele b (A). Positive selection works when the average fitness of the three genotypes genotype (shown in black) is in the order of b/b < B/b < B/B (B, C), while negative selection works when the order is b/b > B/b > B/B (D, E). Balancing selection works such that the average fitness is in the following order: b/b < B/b > B/B (F).

The purpose of this work is to theoretically understand how often turnover of a new sex-determining A/a locus occurs in a more general situation without assuming balancing selection at the linked B/b locus. We here mathematically describe the probability that a newly arisen sex-determining locus turns over the old one, which is referred to as the establishment probability. We found that the establishment probability is markedly high in the case of balancing selection, while it is very low in other modes of selection. It is indicated that the mode of selection at the B/b locus is critical in determining the fate of a newly arisen sex-determining allele. Therefore, to understand the rate of turnover, it is necessary to know how many sexually antagonistic loci are under balancing selection. Our simulations provide insight into how to distinguish the mode of selection from the pattern of polymorphism around the sex-determining locus.

## Model

We use a discrete-generation model of a diploid species with population size *N*. Three loci are considered in the model (Figure 1A). One is the ancestral sex-determining locus, which is located on the sex chromosome where males have genotype XY and females have genotype XX. Although male heterogamety (XY system) is assumed here, our model can also handle female heterogamety (ZW system) by swapping males and females. The model includes another sex-determining locus A/a on the autosome, at which the initial state is that allele a is fixed, so that the locus does not play a role in sex determination. Then, we consider that masculinizing or feminizing allele A just arises by mutation at locus A/a. We are interested in how this new sex-determining allele A behaves in a finite population and how often it spreads and eventually enables the A/a system to override the old X/Y system.

In our model, if allele A has a masculinizing effect, its turnover does not change the heterogamety sex and involves four genotypes (Case 1, Figure 1B). If allele A has a *strongly* feminizing effect (i.e., strong enough to make XYAa a female) (van Doorn and Kirkpatrick 2010), the turnover changes the heterogamety system and involves six genotypes (Case 2, Figure 1C). In either case, to evaluate the effect of the B/b locus, we assume that the levels of fitness of the new and old systems are identical. Therefore, the fate of allele A is mainly determined by the selection effect of another linked locus B/b. It is assumed that the recombination rate is *r* between the A/a and the B/b loci. We ignore mutation at the A/a locus to trace the fate of a single mutation, whereas at the B/b locus, recurrent mutation is allowed between alleles B and b such that this locus should be under selection-mutation balance. The mutation rate from allele B to allele b is assumed to be *u* and *v* denotes the reverse mutation rate from allele b to allele B. The frequency of allele B is denoted by *p*. The fitness of genotypes BB, Bb, and bb is given by 1 + *s*_*m*_, 1 + *h*_*m*_*s*_*m*_, and 1 in males and 1 + *s* _*f*_, 1 + *h*_*f*_ *s* _*f*_, and 1 in females, respectively. As we assume allele B is beneficial for allele A, we set *s*_*m*_ > 0 and *s* _*f*_ < 0 when allele A has a masculinizing effect (Case 1), and *s*_*m*_ < 0 and *s* _*f*_ > 0 is set when allele A has a feminizing effect (Case 2).

### Data Availability

The authors state that all data necessary for confirming the conclusions presented in the manuscript are represented fully within the manuscript. Codes used for numerical analyses and simulation experiments is available at https://github.com/TSakamoto-evo/sex_codes_2020.

## Results

We derive the probability that a single sex-determining mutation at the A/a locus spreads in the population so that the A/a locus becomes the new sex-determining locus and the old X/Y locus becomes monomorphic under the three-locus model illustrated in Figure 1A. This probability is essentially identical to the probability that the new sex-determining mutation successfully increases in frequency by avoiding extinction immediately after its introduction, as pointed out by van Doorn and Kirkpatrick (2007, 2010). Once the frequency of the new sex-determining mutation increases, it is very likely that it further increases to an intermediate frequency so that the sex-determining locus transitions from the X/Y locus to the A/a locus. This is because our model assumes that the new A/a system together with the linked B/b sex antagonistic locus has higher fitness than the old X/Y system. In the following, we derive this probability of the successful spread of allele A, which is referred to as the “establishment probability.” The “establishment” means that the new system is used by all individuals in the population (i.e., the new system is fixed but alleles A and a coexist stably), rather than the old and new systems coexisting in the population. The establishment probability is derived separately for the two cases (Cases 1 and 2).

### Case 1: Turnover without changing the heterogametic sex

We first consider Case 1, where allele A has a masculinizing effect (Figure 1B). For allele B to be beneficial for allele A, we assume *s*_*m*_ > 0 and *s* _*f*_ < 0 in this section. We derive the probability of allele A escaping immediate extinction by using the branching process. Let *φ*_*i*_(*p*_0_) (*i* ∈ {*B, b*}) be the establishment probability of allele A that arises in linkage with allele *i* at the B/b locus when the frequency of allele B is *p*_0_. If allele A links with male-beneficial allele B, allele A is favored by linked selection. On the other hand, if allele A links with female-beneficial allele b, allele A is disfavored by linked selection. By denoting the frequencies of haplotypes A-B and A-b by *x*_*B*_ and *x*_*b*_, respectively, and ignoring second-order terms in *s*_*m*_, *r, u* and *v*, the expected changes of the frequencies of A-B andA-b in one generation are given by

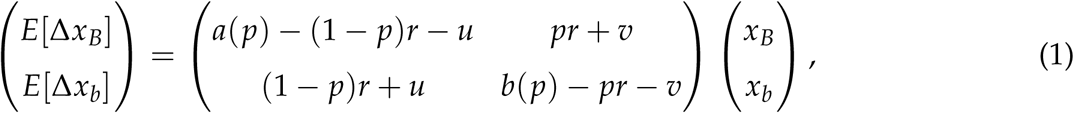

where *a*(*p*) = *h*_*m*_*s*_*m*_ + (*s*_*m*_ − 3*h*_*m*_*s*_*m*_)*p* + (2*h*_*m*_*s*_*m*_ − *s*_*m*_)*p*^2^ and *b*(*p*) = −*h*_*m*_*s*_*m*_ *p* + (2*h*_*m*_*s*_*m*_ − *s*_*m*_)*p*^2^ (for details, see APPENDIX B). It is interesting to point out that Equation 1 involves only selection parameters in males, not those in females. This means that *E*[Δ*x*_*i*_] directly depends on selection among males, but does not necessarily mean that selection does not work in females. Selection in females is involved because it affects *p*, the frequency of allele B.

In the following, we first derive the establishment probability for a special case where selection at the B/b locus does not provide any systematic pressure on *p*. In this case, *p* does not change rapidly, so we can treat it as a constant, at least in the timescale of a newly arisen allele escaping from initial extinction (see APPENDIX C). With this assumption, we can obtain the establishment probability as a solution of a cubic equation, which is given by a function of *p*_0_, the frequency of allele B when allele A arises. Next, we consider a more general case where *p* changes, from the initial value *p*_0_ to the equilibrium value *p*\*. In this case, by contrast, we show that the establishment probability is given by a function of both *p*_0_ and *p*\*, of which the effect of *p*\*is quite large. In either case, we first obtain the establishment probability conditional on *p*_0_, and then we derive the unconditional establishment probability by incorporating the stationary distribution of *p*_0_.

### Establishment probability when a constant p can be assumed

We consider a case where we can assume *p* does not change significantly in the timescale in which we are interested in. In other words, the expected change of *p* in one generation, *M*_*p*_, is assumed to be as small as 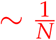, where *M*_*p*_ is given by:

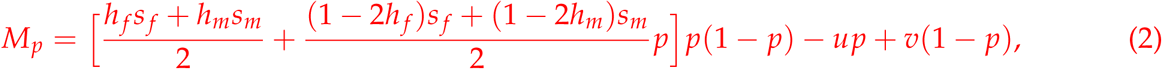

if the second-order terms of *s*_*m*_, *s* _*f*_, *u*, and *v* are ignored. This assumption holds when *h*_*m*_ ≈ *h*_*f*_ and *s*_*m*_ + *s*_*f*_ ≈ 0, so that the levels of selection in the two sexes are of the same strength, but work in opposite directions. This mode of selection is referred to as “equalizing selection”. We assume that *s*_*m*_, *s* _*f*_, *u, v*, and *r* are so small that their second-order terms can be ignored. Then, following the branching process approximation (Barton 1987), it is straightforward to show that the two establishment probabilities satisfy:

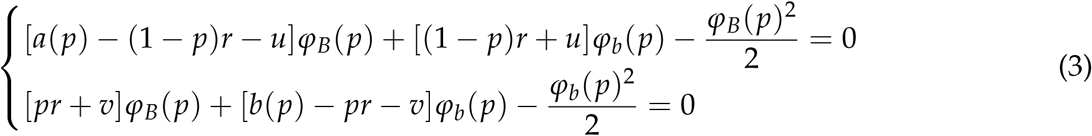

(for details, see APPENDIX C). Notably, similar models were used to analyze the effect of a linked allele on the establishment of locally beneficial alleles (Aeschbacher and Bürger 2014) or the effect of population structure on the establishment of a beneficial mutation (Pollak 1966; Barton 1987; Tomasini and Peischl 2018; Sakamoto and Innan 2019). Equations 3 can be reduced to a cubic equation and be solved analytically (for details, see APPENDIX D). It is important to note that establishment is promoted by selection if *φ*_*B*_(*p*) is positive; otherwise, allele A is likely selected against and goes extinct. Whether *φ*_*B*_(*p*) > 0 depends on the leading eigenvalue of the matrix in Equation 1. When the leading eigenvalue is positive, *φ*_*B*_(*p*) and *φ*_*b*_(*p*) are given by:

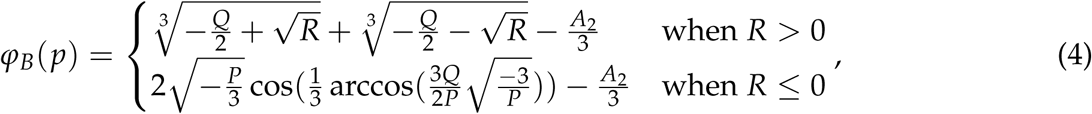

and

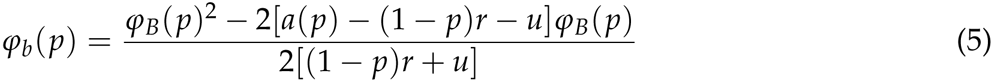

where

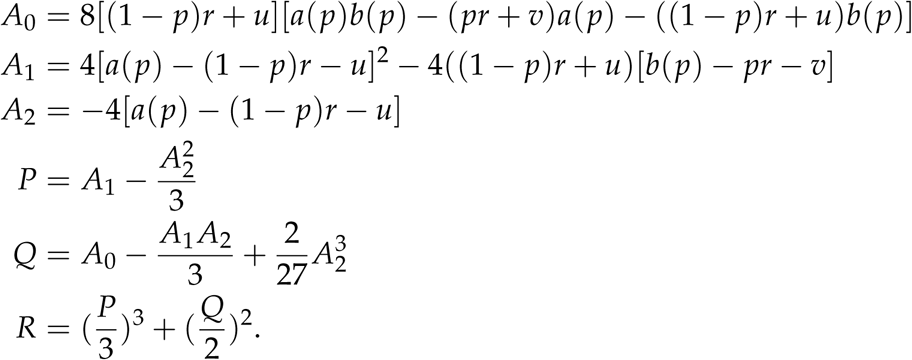

When *r* is small, they can be approximated in quite simple equations:

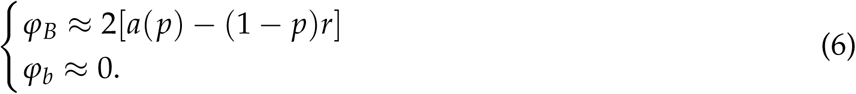

We performed simple forward simulations in a Wright–Fisher population to check the accuracy of our derivation (for details of the simulations, see APPENDIX A). We confirmed that Equation 3 is in excellent agreement with the simulations for a wide range of the parameters. Some of the results are shown in Figure 3, where the two establishment probabilities, *φ*_*B*_(*p*_0_) and *φ*_*b*_(*p*_0_), are plotted by assuming *s*_*m*_ = −*s*_*f*_ = 0.02, *N* = 10, 000. Through this work, the mutation rates at locus B/b are fixed to be quite small values, *u* = *v* = 1.0 × 10^−6^, unless otherwise mentioned. Note that the effect of mutation rate is small in the establishment process unless the mutation rate is very large (but the mutation matters when the stationary distribution of allele B is considered, as demonstrated below).

**Figure 3.**
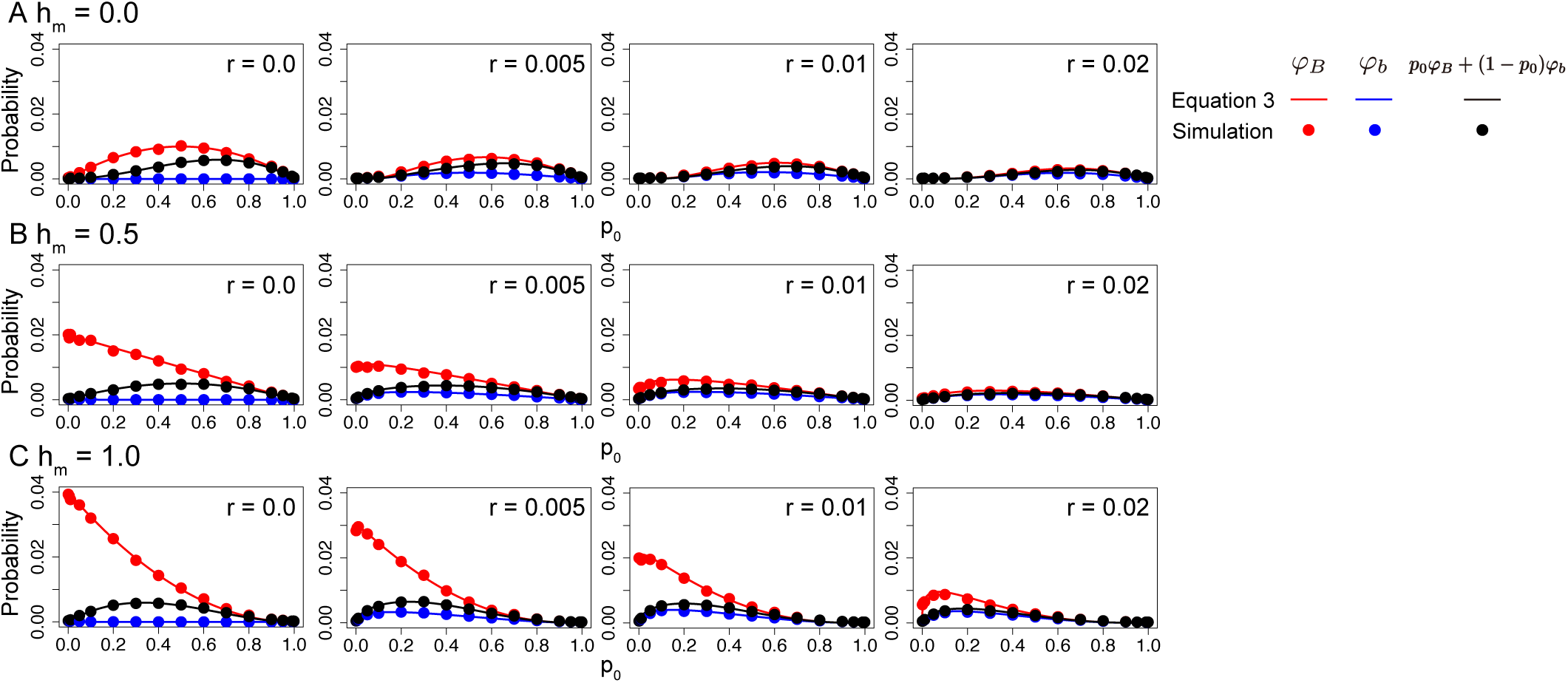
Establishment probability of allele A for different dominance and recombination rates. Three dominance coefficients are assumed: (A) *h*_*m*_ = 0.0, (B) *h*_*m*_ = 0.5 and (C) *h*_*m*_ = 1.0. The other parameters are as follows: *s*_*m*_ = − *s* _*f*_ = 0.02, *h*_*f*_ = *h*_*m*_, *N* = 10, 000, *u* = *v* = 1.0 × 10^−6^. Error bars on the red and blue circles represent the 95 % confidence interval, but they are too small to be seen.

We first focus on the case of *r* = 0, that is, the A/a and B/b loci are completely linked, in order to investigate the effect of dominance (*h*). Suppose that allele B is recessive (*h* = 0, left panel in Figure 3A). Then, a newly arisen allele A can benefit from the B/b locus only when it arises in a BB homozygote. In such a case, *φ*_*B*_ increases as *p*_0_ increases to *p*_0_ = 0.5 (plotted in red), where the effect of sexually antagonistic selection is maximized. *φ*_*B*_ decreases as *p*_0_ decreases from 0.5 to 1, making a symmetric function. With the assumption of no recombination, it is obvious that *φ*_*b*_ ≈ 0 for any *p*_0_ (plotted in blue). The weighted average of *φ*_*B*_ and *φ*_*b*_ (i.e., *p*_0_ *φ*_*B*_ + (1 − *p*_0_)*φ*_*b*_) is plotted in black.

In contrast, in the dominant case (*h* = 1, left panel in Figure 3C), *φ*_*B*_ for a small *p*_0_ is quite high because a newly arisen allele A in linkage with allele B is immediately selected for, regardless of the genotype at the B/b locus. This selection works particularly efficiently when *B* is so rare that the selective advantage of A-B haplotypes is large in comparison with the population fitness. Therefore, *φ*_*B*_ is given by a monotonically increasing function with decreasing *p*_0_, but as *p*_0_ decreases the probability that allele A arises in linkage with allele B decreases; therefore, the weighted average has a peak in the middle. An intermediate pattern is observed in a case of partial dominance (*h* = 0.5, left panel in Figure 3B). Similar results were obtained for other values of selection intensity as long as *s*_*m*_ = − *s* _*f*_ (not shown).

We next consider the effect of recombination. The approximations given by Equation 6 agree overall with the simulation results as long as *r* is small. As the recombination rate increases, *φ*_*B*_ decreases because the association with allele B becomes weaker. When *r* is relatively small, *φ*_*b*_ increases as the recombination rate increases because recombination gives a chance to link with allele B and the A-B association may be preserved if further frequent recombination does not break the association. However, this benefit does not hold for a large *r*, and *φ*_*b*_ decreases as the recombination rate increases because further recombination prevents the stable linkage of allele A and allele B and reduces the benefit of linkage.

### Establishment probability when p changes

We next consider the case where *p* can change during the establishment process due to selection and mutation (i.e.,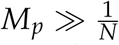). To derive *φ*_*B*_(*p*_0_) and *φ*_*b*_(*p*_0_), we use a continuous time approximation of the branching process (Barton 1995). With a deterministic approximation for *M*_*p*_ (*M*_*p*_ ≠ 0), the two establishment probabilities satisfy:

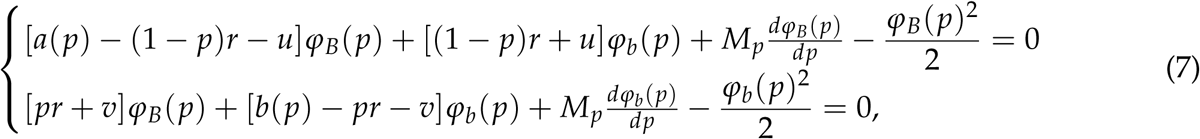

(for details, see APPENDIX C). This differential equation can be solved numerically by setting an initial condition, for which we use the establishment probability of allele A that arises when *p* is at stable equilibrium *p*\*. *φ*_*B*_(*p*\*) and *φ*_*b*_(*p*\*) can be numerically computed by Equation 3 because we can assume *p* does not change significantly around *p* *. Technically, we cannot use the exact values of *φ*_*B*_(*p*\*) and *φ*_*b*_(*p*\*) as an initial condition because they violate the assumption of *M*_*p*_ ≠ 0. To avoid this problem, assuming a very small *ε*, we use *φ*_*B*_(*p*\*± *ε*) ≈ *φ*_*B*_(*p*\*) and *φ*_*b*_(*p*\*± *ε*) ≈ *φ*_*b*_(*p*\*) as an initial condition, which does not markedly affect the numerical solutions as long as *ε* is small.

When *p* is not constant, both *p*_0_ and *p*\*play an important role in determining *φ*_*B*_(*p*_0_) and *φ*_*b*_(*p*_0_). The relative contribution of *p*\*can be large when selection is strong and *p*_0_ is not far from *p*\*, so that *p* quickly approaches *p*\*. In such a case, *φ*_*B*_(*p*_0_) and *φ*_*b*_(*p*_0_) cannot be large when *φ*_*B*_(*p*\*) and *φ*_*b*_(*p*\*) are very small. This argument can be explained as follows. Let us consider the fate of the descendants of an allele A that arises. After the mutation arises, *p* changes and finally reaches around *p*\*. Denote the numbers of haplotypes A-B and A-b when *p* reaches *p*\*as *X*_*B*_ and *X*_*b*_, respectively. Then, the establishment probability is approximately given by:

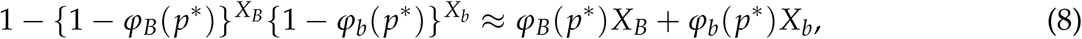

unless the population is very small.

Because *p*\*largely depends on the mode of selection on the B/b locus (see Figure 2), we here consider *φ*_*B*_(*p*_0_) and *φ*_*b*_(*p*_0_) under three modes of selection separately (i.e., balancing, negative, and positive selection on allele B). For each mode of selection, we performed extensive forward simulations to check the performance of Equation 7. It was found that *φ*_*B*_(*p*_0_) and *φ*_*b*_(*p*_0_) computed by Equation 7 well agreed with the simulation results for a wide range of parameter space, and representative cases are shown in Figures 4, 5 and 6.

**Figure 4.**
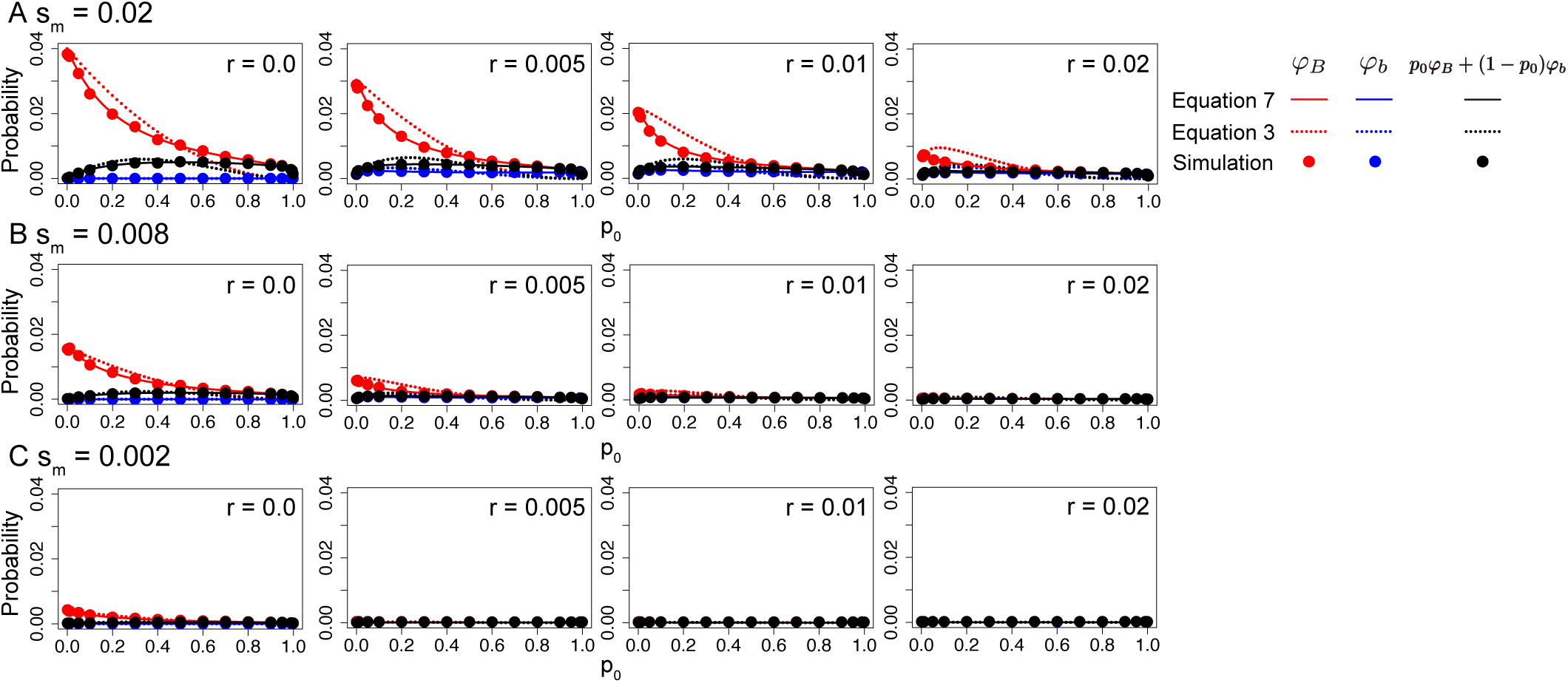
Establishment probability for the case of balancing selection on allele B. Different strengths of selection are assumed: (A) *s*_*m*_ = −*s*_*f*_ = 0.02, (B) *s*_*m*_ = −*s*_*f*_ = 0.008 and (C) *s*_*m*_ = −*s*_*f*_ = 0.002. Other parameters are assumed to be *h*_*m*_ = 1.0, *h*_*f*_ = 0.0, *N* = 10, 000, *u* = *v* = 1.0 × 10^−6^. Error bars on the red and blue circles represent the 95 % confidence interval, but they are too small to be seen.

**Figure 5.**
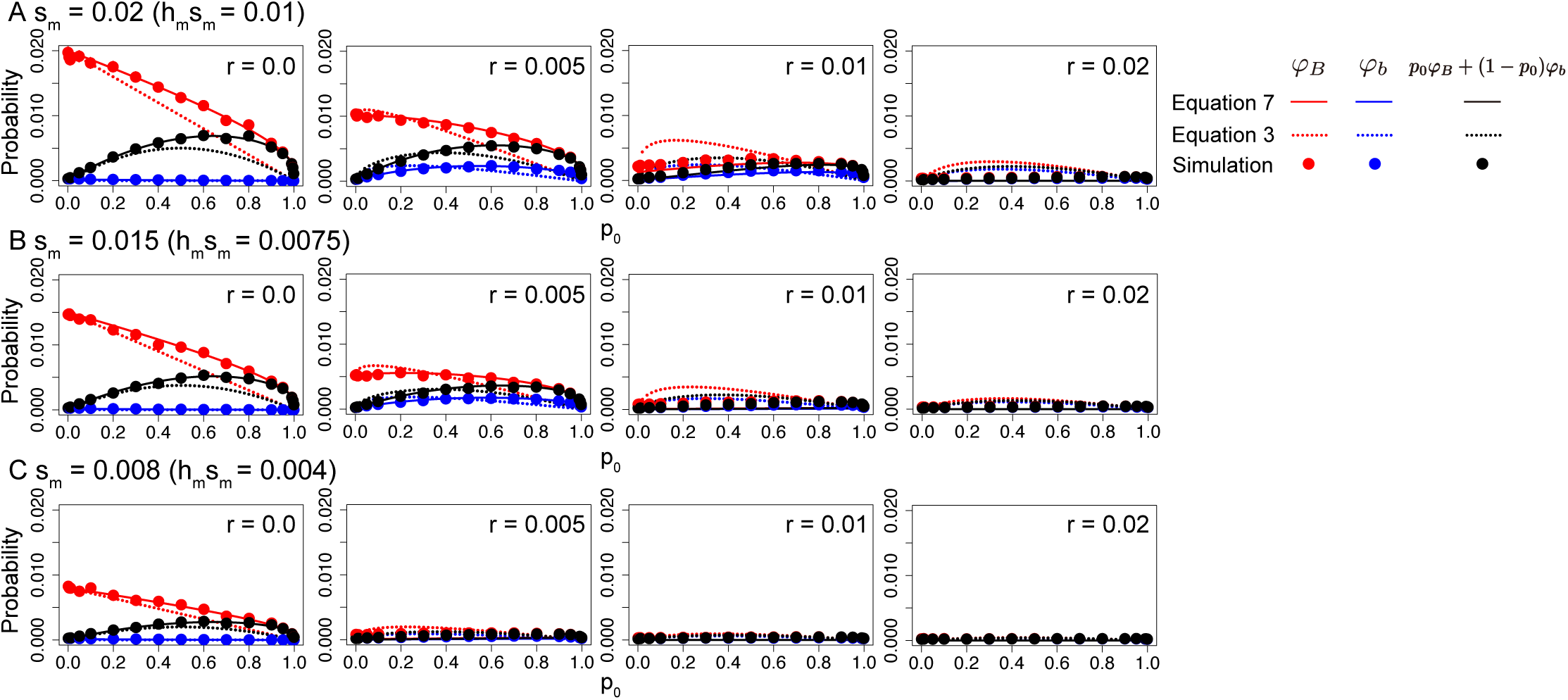
Establishment probability for the case of negative selection against allele B. Different strengths of selection are assumed: (A) *s*_*m*_ = 0.02, (B) *s*_*m*_ = 0.015 and (C) *s*_*m*_ = 0.008. Other parameters are assumed to be *s* _*f*_ = −2*s*_*m*_, *h*_*m*_ = *h*_*f*_ = 0.5, *N* = 10, 000, and *u* = *v* = 1.0 × 10^−6^. Error bars on the red and blue circles represent the 95 % confidence interval, but they are too small to be seen.

**Figure 6.**
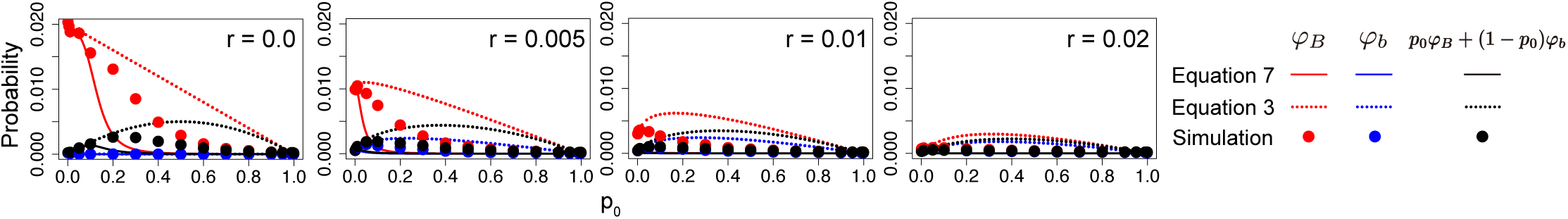
Establishment probability for the case of positive selection for allele B. Other parameters are assumed to be *s*_*m*_ = 0.02, *s* _*f*_ = − 0.01, *h*_*m*_ = *h*_*f*_ = 0.5, *N* = 10, 000, and *u* = *v* = 1.0 × 10^−6^. Error bars on the red and blue circles represent the 95 % confidence interval, but they are too small to be seen.

First, we consider the case of balancing selection. To make a realization of balancing selection, it is necessary to set *h*_*m*_ large enough to secure the average fitness of B/b heterozygote higher than 1. When *h*_*m*_ is large, a newly arisen allele A would be immediately selected for together with allele B. Therefore, the overall behavior of *φ*_*B*_(*p*_0_) and *φ*_*b*_(*p*_0_) could be quite similar to that in the case of a high *h* when *p* does not change (e.g., Figure 3C where *h*_*m*_ = 1 is assumed). This is demonstrated in Figure 4A, where we assume *s*_*m*_ = 0.02, *s* _*f*_ = − 0.02, *h*_*m*_ = 1.0, *h*_*f*_ = 0.0 (as a consequence, *p*\*= 0.5), while all other parameters are the same as those used in Figure 3. To emphasize the difference from the case of a constant *p*, this figure also shows *φ*_*B*_(*p*_0_) and *φ*_*b*_(*p*_0_) computed by Equation 3 for comparison. The major difference from Equation 3 is that, when *p*_0_ *< p*\*, *φ*_*B*_(*p*_0_) is lower than that in the case of constant *p* (the red broken lines in Figure 4), because the selective advantage of haplotype A-B would be reduced by the rapid increase in the frequency of allele B by selection (i.e., both haplotypes A-B and a-B increase). *φ*_*b*_(*p*_0_) is overall very low, similar to the case of a constant *p* (Figure 3C). On the other hand, when *p*_0_ *> p*\*, *φ*_*B*_(*p*_0_) is larger than that in the case of constant *p* because the number of males having allele B decreases, with which haplotype A-B has to compete. Figures 4B and C are for weaker selection coefficients (*s*_*m*_ = − *s* _*f*_ = 0.008 and 0.002), where *φ*_*B*_(*p*_0_) and *φ*_*b*_(*p*_0_) are overall reduced and the difference from Equation 3 is small perhaps because *p* does not change rapidly with weak selection.

We next consider the case of negative selection, where *p* would move from *p*_0_ to *p*\*, which is usually very low. If a very low *p*\*is assumed, from Equation 3, we can approximate *φ*_*B*_(*p*\*) and *φ*_*b*_(*p*\*) as:

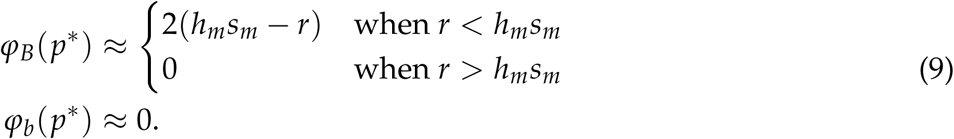

These equations mean that the establishment of allele A is very unlikely when *r > h*_*m*_*s*_*m*_. Thus, the major difference from the case of a constant *p* is that, as the recombination rate increases, the establishment rate decreases to ∼ 0 around the threshold *r > h*_*m*_*s*_*m*_ (see Appendix E for the behavior for large *r*). This is demonstrated in Figure 5A, where *s*_*m*_ = 0.02, *s* _*f*_ = −0.04, *h*_*m*_ = *h*_*f*_ = 0.5 are assumed such that the comparable result for the case of a constant *p* is Figure 3B with the same selection parameters for males (all other parameters are identical). In Figure 5A, *r* = *h*_*m*_*s*_*m*_ holds at *r* = 0.01, which works as the threshold. When *r* is smaller than this threshold (*r* = 0.0 and 0.005 in Figure 5A), *φ*_*B*_(*p*_0_) and *φ*_*b*_(*p*_0_) are roughly in agreement with those in Figure 3B, although *φ*_*B*_(*p*_0_) and *φ*_*b*_(*p*_0_) are slightly higher than the case of a constant *p*, especially when *p*_0_ is not small. The situation changes dramatically as the recombination rate exceeds the threshold (i.e., *r* = 0.02 in Figure 5B): *φ*_*B*_(*p*_0_) and *φ*_*b*_(*p*_0_) decrease to as low as ∼ 2/*N* (i.e., the neutral expectation for a sex-determining allele), where there is no benefit of linked selection and only drift-driven establishment occurs in a nearly neutral fashion. In contrast, in Figure 3B, *φ*_*B*_(*p*_0_), *φ*_*b*_(*p*_0_) ≫ 2/*N* unless *p*_0_ is close to 0 or 1. Such strong reduction of *φ*_*B*_(*p*_0_) when *r* exceeds *h*_*m*_*s*_*m*_ (Figure 5A) can be explained as follows. A newly arisen allele A benefits from linked selection when it arises in association with allele B. Once the linkage is broken by recombination, allele A has almost no chance to recombine back to link to allele B because the frequency of allele B is very low. Therefore, after allele A loses linkage with allele B, there would be no selection for allele A so that the establishment of allele A has to rely on random genetic drift. Figures 5B and C show the results for weaker selection (*s*_*m*_ = 0.015 and 0.008), where the general pattern is similar to that in Figure 5A, while *φ*_*B*_(*p*_0_) and *φ*_*b*_(*p*_0_) are overall reduced.

We finally consider the case of positive selection, where *p*\* is generally very large (i.e., ≈ 1). Unless *p*_0_ is small, *p* increases very quickly to *p*\* ≈ 1, that is, the B/b locus is almost fixed for allele B. Therefore, even when allele A arises in association with allele B, allele A does not benefit from linked selection on locus B, resulting in a very low *φ*_*B*_(*p*_0_). Particularly when allele A at *p* ≈ *p*\*, random genetic drift plays a role in the establishment process. The theoretical treatment for this situation is shown in APPENDIX E. Figure 6 shows *φ*_*B*_(*p*_0_) and *φ*_*b*_(*p*_0_) for the case of positive selection assuming *s*_*m*_ = 0.02, *s* _*f*_ = − 0.01, *h*_*m*_ = *h*_*f*_ = 0.5. It is demonstrated that allele A can spread efficiently with the help of linked selection for allele B only when *p*_0_ is small and *r* is so small that the initial association between A and B can be maintained for a while. The performance of Equation 7 is not as good as that in the cases of balancing selection and negative selection. It appears that Equation 7 underestimates the establishment probability because our derivation based on the branching process ignores establishment events occurring in a nearly neutral fashion.

### Unconditional establishment probability

In the above, we consider the establishment probability as a function of the initial frequency of allele B, *p*_0_. We are here interested in the unconditional establishment probability, which is the weighed average over the stationary distribution of *p*_0_. Following Wright’s formula (Wright 1931), the stationary distribution of *p*_0_, *g*(*p*), is given by:

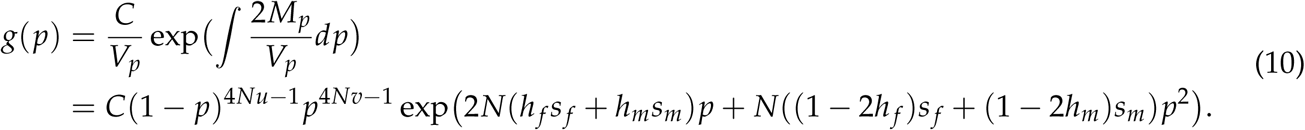

where 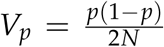 and *C* is determined such that 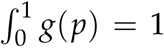 (Connallon and Clark 2012). It is well recognized that this formula works very well when the selection intensities, *s* _*f*_ and *s*_*m*_, are relatively small so that their second-order terms are negligible (Crow and Kimura 1970; Ewens 2004). An intriguing exceptional case is when selection is weak but the absolute values of the selection intensities are not small. In the previous section, we considered a case where selection in males and that in females are well balanced when *h*_*f*_ ≈ *h*_*m*_ and *s* _*f*_ + *s*_*m*_ ≈ 0, irrespective of the absolute values of *s* _*f*_ and *s*_*m*_. In such a special case of equalizing selection, the stationary distribution may be better given by:

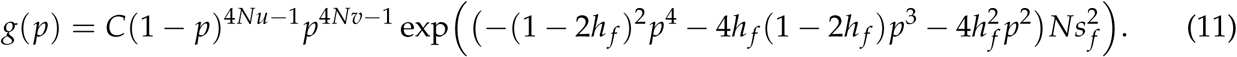

Then, the unconditional establishment probability of allele A, *φ*, is given by:

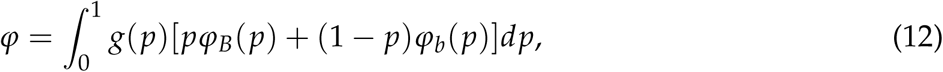

where *φ*_*B*_(*p*) and *φ*_*b*_(*p*) are given by Equation 3 for *h*_*f*_ ≈ *h*_*m*_ and *s* _*f*_ + *s*_*m*_ ≈ 0, and otherwise by Equation 7.

We can obtain an approximation of the establishment probability in a simple form for several special cases of *r* = 0. When sexually antagonistic selection works as balancing selection, the establishment probability for *r* = 0 is approximated by:

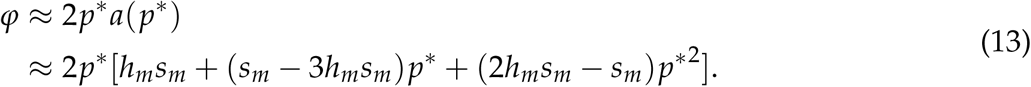

It is indicated that *φ* is on the same order of magnitude as *s*_*m*_.

In the case of negative selection, assuming that selection is much stronger than mutation (*u* and *v*) and *h*_*m*_ is not very small, we have *p*\* ≈ 0 and *φ* ∼ *φ*_*b*_(0). Furthermore, because *φ*_*B*_(*p*) ≫ *φ*_*b*_(*p*) and *M*_*p*_ ≈ 0, Equation 7 for *r* = 0 is roughly simplified:

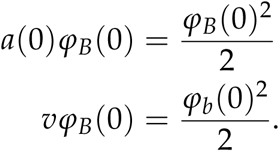

Under these approximations, *φ* is given by

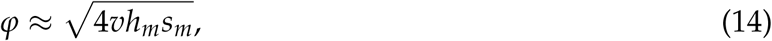

indicating that *φ* is on the order of 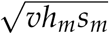. For selection to be dominant over random genetic drift, *N*^2^*vh*_*m*_*s*_*m*_ ≫ 1 is required.

If sexually antagonistic selection works as positive selection and the mutation rate is low, linked selection no longer works and establishment is driven by random genetic drift, as discussed above. Then, the establishment probability is roughly given by 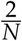. In such cases, sexually antagonistic selection does not markedly increase the establishment probability.

The unconditional establishment probability computed by Equation 12 is plotted for the four modes of sexually antagonistic selection in Figure 7. The approximations for *r* = 0 (Equations 13, 14) are also presented by triangles on the y-axis, which show excellent agreement with the exact formula and simulation results. Three different values of the mutation rate are considered (*u* = *v* = 10^−5^, 10^−6^, 10^−7^).

**Figure 7.**
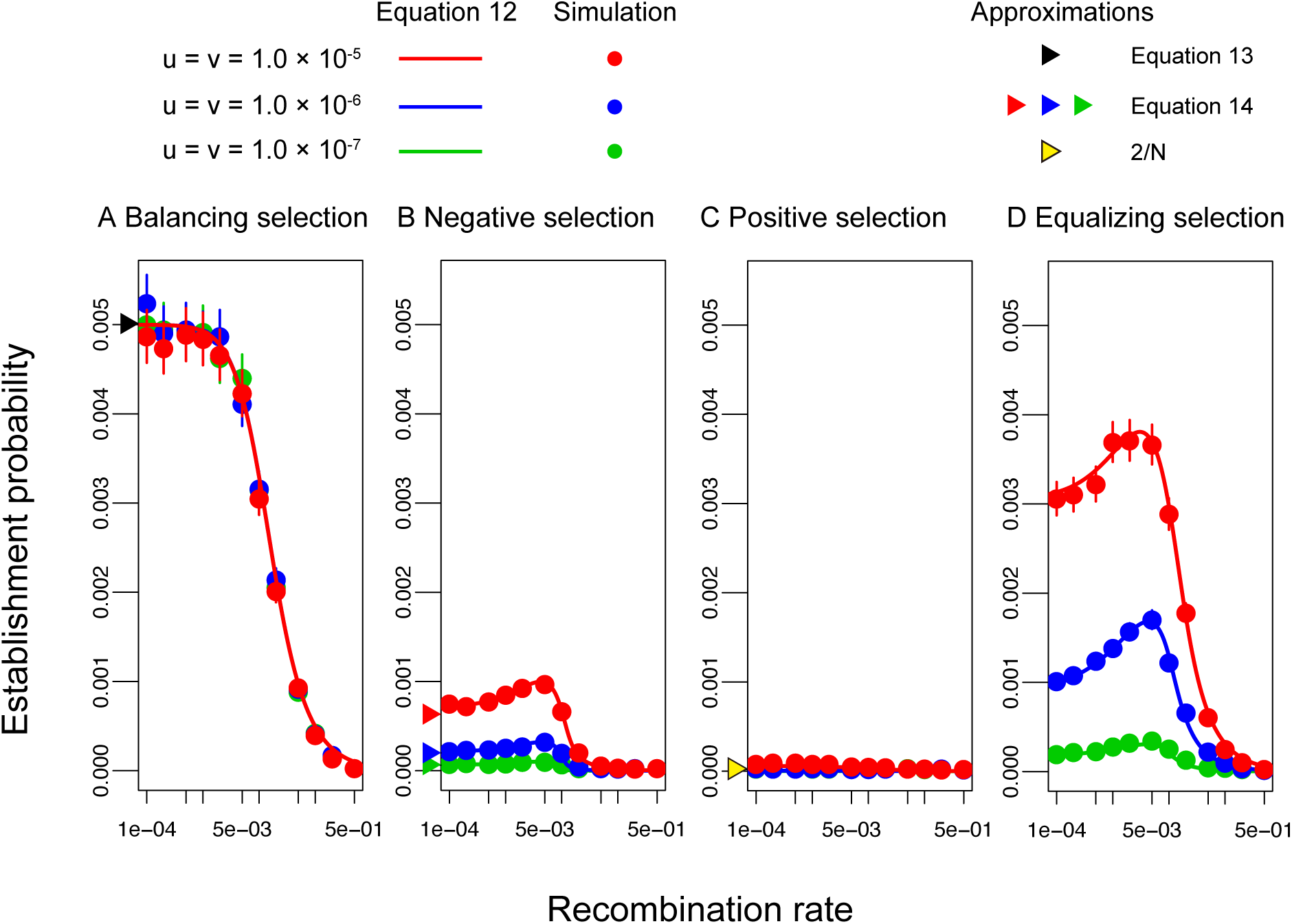
Establishment probability of a masculinizing allele for different modes of sexually antagonistic selection. *N* = 100, 000 and *u* = *v* are assumed. Other parameters are (A) *s*_*m*_ = 0.02, *s* _*f*_ = − 0.02, *h*_*m*_ = 1.0, *h*_*f*_ = 0.0, (B) *s*_*m*_ = − 0.02, *s* _*f*_ = 0.025, *h*_*m*_ = *h*_*f*_ = 0.5, (C) *s*_*m*_ = 0.02, *s* _*f*_ = − 0.01, *h*_*m*_ = *h*_*f*_ = 0.5 and (D) *s*_*m*_ = 0.02, *s* _*f*_ = − 0.02, *h*_*m*_ = *h*_*f*_ = 0.5. Similar results were obtained for *N* = 10, 000. Error bars for circles represent the 95 % confidence interval.

The establishment probability is in general highest when balancing selection works (Figure 7A). This is because an intermediate *p* provides both the benefit of linkage and a higher chance to link with allele B to allele A. Because the mutation rate at locus B/b does not markedly influence the stationary distribution of *p*, the establishment probability does not depend on the mutation rate either.

When negative selection works at locus B, the establishment probability is quite low (Figure 7B) because negative selection keeps allele B very rare. Mutation from allele b to B contributes to the establishment of allele A in two ways. First, it increases the frequency of beneficial allele B, resulting in a higher chance that allele A acquires linkage with allele B by recombination. Second, it increases the probability that haplotype A-b mutates to A-B. As a consequence, as the mutation rate *v* increases, the establishment probability becomes larger.

When positive selection works, the establishment probability is very low for all three mutation rates (Figure 7C). This is because allele B is already fixed and linkage with allele B is no longer beneficial. Allele A may benefit from linked selection only when *u* is extremely high.

Thus, if selection is directional, *p*\* is very low or high (under negative or positive selection, respectively), which does not markedly help the establishment of allele A. On the other hand, if balancing selection maintains alleles B and b in an intermediate frequency, allele A can take advantage of it. An intermediate situation is when equalizing selection works ((Figure 7D), where the process is nearly neutral because *s* _*f*_ + *s*_*m*_ ≈ 0 and *h*_*f*_ ≈ *h*_*m*_). Because *p* fluctuates between 0 and 1 by random genetic drift, allele A can likely benefit from locus B/b if it arises when *p* is intermediate. This is why the establishment probability is largely affected by the mutation rate at locus B/b, which determines how often mutation is produced.

It is interesting to note that very tight linkage does not necessarily increase the unconditional establishment probability (e.g., Figures 7B, D). With an increasing the recombination rate, in general, *φ*_*b*_ increases while *φ*_*B*_ decreases because recombination enhances exchange between haplotype A-B and haplotype A-b. Because the unconditional probability is given by the weighted average of *φ*_*B*_ and *φ*_*b*_, there appears to be an optimal recombination rate to maximize *φ*.

### The pattern of neutral polymorphism after turnover

To investigate the effect of the mode of selection at locus B/b on the pattern of neutral polymorphism in a surrounding region, we performed forward simulations under the infinite-sites model. The spatial distributions of the nucleotide diversity after turnover in a typical run are plotted for each of the four modes of selection in Figure 8. The amount of nucleotide variation is measured by the average pairwise differences (*π*) standardized by the neutral expectation (*θ* = 4*Nµ* where *µ* is the neutral mutation rate). Haplotype *i* (*i* ∈{A, a}) is denoted as the haplotype with allele *i*, and Figure 8 plots three *π* values: *π*_*A*−*A*_ for *π* within haplotype A (red line); *π*_*a*−*a*_ for *π* within haplotype a (blue line); and *π*_*A*−*a*_ for *π* between haplotypes

**Figure 8.**
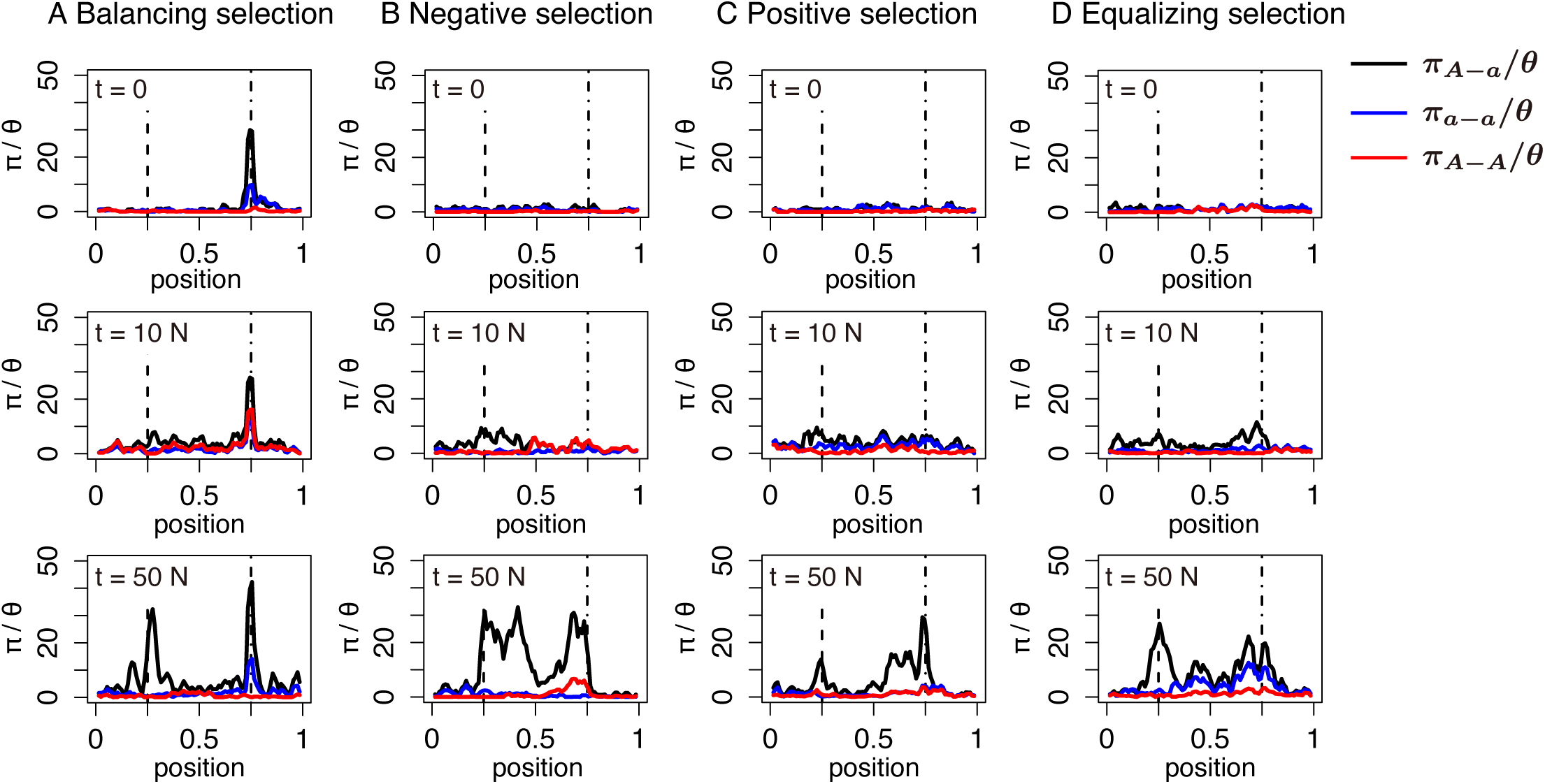
The temporal dynamics of nucleotide diversity after turnover of the sex-determining locus. Results for a single simulation run with *N* = 1, 000 are shown for each mode of selection. The spacial distributions of the normalized *π*_*A*−*A*_, *π*_*a*−*a*_ and *π*_*A*−*a*_ are shown in red, blue, and black, respectively. In the simulation, locus A/a is located at relative position 0.25 (vertical dashed lines) in the simulated region, while locus B/b is located at relative position 0.75 (vertical dot-dashed lines). *t* is denoted as the number of generations since the new sex-determination system was fixed (i.e., since the old sex-determining locus X/Y became monomorphic). The recombination rate of the entire region was assumed to be 0.002 such that the recombination rate between loci A/a and B/b = 0.001. The selection parameters used for each mode of selection were (A) *s*_*m*_ = −*s*_*f*_ = 0.02, *h*_*m*_ = 1.0, *h*_*f*_ = 0.0, (B) *s*_*m*_ = 0.02, *s* _*f*_ = −0.04, *h*_*m*_ = *h*_*f*_ = 0.5, (C) *s*_*m*_ = 0.025, *s* _*f*_ = −0.02, *h*_*m*_ = *h*_*f*_ = 0.5 and (D) *s*_*m*_ = −*s*_*f*_ = 0.02, *h*_*m*_ = *h*_*f*_ = 0.5.

A and a (black line). Let us first focus on the case of *t* = 0 (i.e., just after the turnover has completed). When balancing selection works, *π*_*A*−*a*_ has a striking peak around the locus B/b (Figure 8A), and a weaker peak is observed for *π*_*A*−*A*_ and *π*_*a*−*a*_ due to recombination between the two loci. This is because the B/b polymorphism had been maintained for a very long time by balancing selection before the turnover occurred, which is not observed in the other modes of selection (Figures 8B-D).

As time passes, in all four cases, neutral mutations start to accumulate around the A/a locus, making a novel peak of divergence between haplotypes A and a (i.e., *π*_*A − a*_). The growth of the peak at the A/a locus proceeds while maintaining the peak at the B/b locus in the case of balancing selection (Figure 8A). The pattern is as predicted by Kirkpatrick and Guerrero (2014). In contrast, in the other three cases (Figures 8B-D), a peak newly arises at the B/b locus and grows along with the peak at the A/a locus. Thus, the patterns are similar in the cases of negative, positive and equalizing selection, whereas balancing selection is unique in that the peak at the B/b locus already exists, which is much higher than that at the A/a locus shortly after the turnover.

### Case 2: Turnover with changing heterogametic sex

We next consider Case 2, where allele A has a feminizing effect so that the heterogametic sex changes from male to female. That is, Case 2 assumes *s* _*f*_ > 0 for allele B to be beneficial for allele A. As in Case 1, allele A can spread if allele A successfully avoids extinction shortly after it arises (van Doorn and Kirkpatrick 2010). Therefore, we again use the approximation of the branching process, following Case 1. By ignoring the second-order terms in *s* _*f*_, *r, u*, and *v*, the expected change of the frequency in one generation is given by:

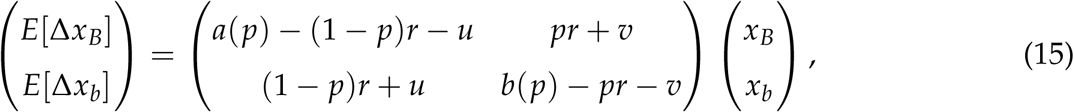

where *a*(*p*) = *h*_*f*_ *s* _*f*_ + (*s* _*f*_ − 3*h*_*f*_ *s* _*f*_)*p* + (2*h*_*f*_ *s* _*f*_ − *s* _*f*_)*p*^2^ and *b*(*p*) = −*h*_*f*_ *s* _*f*_ *p* + (2*h*_*f*_ *s* _*f*_ − *s* _*f*_)*p*^2^ (see APPENDIX B for details).

By comparing Equation 15 with Equation 1, we notice that the two equations are identical if we replace *h*_*f*_ by *h*_*m*_ and *s* _*f*_ by *s*_*m*_. It is indicated that the above arguments for Case 1 are also applicable to Case 2 by changing these variables. When equalizing selection works and *p* does not change rapidly (i.e., *h*_*m*_ ≈ *h*_*f*_ and *s*_*m*_ + *s* _*f*_ ≈ 0), the establishment probability is given by Equation 3. On the other hand, when *p* changes when allele A is rare, the establishment probability depends on both *p*_0_ and stable equilibrium *p*\*. If balancing selection works, the establishment probability is given by Equation 7. When allele B is subject to negative selection, the establishment probability depends on whether *r* is larger or smaller than the threshold that is given by *h*_*f*_ *s* _*f*_. When *h*_*f*_ *s* _*f*_ *> r*, the establishment probability is given by Equation 7. Whereas, when *h*_*f*_ *s* _*f*_ *< r, ϕ*_*B*_(*p*\*) ≈ 0 so that the establishment of allele A would be drift-driven once *p* reaches *p*\* : therefore, *φ*_*B*_(*p*_0_) could be as small as ∼ 1/*N*. When allele B is subject to positive selection, we have *φ*_*B*_(*p*\*) ≈ 0. Consequently, the establishment process is driven by random genetic drift after *p* reaches *p*\*. Therefore, as long as *p*_0_ is near *p*\* and positive selection is strong, the establishment probability is as low as ∼ 1/*N* (for more details, see APPENDIX E). Thus, the process in Case 2 can be well described by the equations developed for Case 1, as is demonstrated in Figures S6-S9. The unconditional establishment probability is also given by Equation 12 (see Figure S10).

## DISCUSSION

In some clades such as teleost fish and amphibians, sex is often determined by a single locus rather than heteromorphic sex chromosomes. In such species, turnover of the sex-determining locus occurs so frequently that genetic divergence around the locus does not proceed. There are many factors that potentially promote the turnover of sex-determining loci, such as random genetic drift (Bull and Charnov 1977; Veller *et al*. 2017; Saunders *et al*. 2018), deleterious mutation load (Blaser *et al*. 2013, 2014), sexually antagonistic selection (van Doorn and Kirkpatrick 2007, 2010), sex ratio bias (Kozielska *et al*. 2010) and haploid selection (Scott *et al*. 2018). Recently, van Doorn and Kirkpatrick (2007, 2010) pointed out that turnover of sex-determining loci could be enhanced by linked selection. That is, a new sex-determining allele (allele A in our model) can be beneficial when it arises near a locus under sexually antagonistic selection (locus B/b). This article is aimed at understanding how often turnover of sex-determining loci occurs with the help of linked selection at a nearby locus. Previous studies mainly by van Doorn and Kirkpatrick (2007, 2010) focused on the deterministic phase to obtain the rate of increase in the frequency of allele A, which is not sufficient to address the question on the rate of turnover, as mentioned in the Introduction. We here provide a full theoretical description of the behavior of allele A from when it newly arises to its establishment, where both the initial stochastic and following deterministic phases are taken into account. We provide some technical comments in APPENDIX F to provide intuitive insights on how to understand the results of van Doorn and Kirkpatrick (2007, 2010) in our framework.

Our theory shows that the establishment probability is given by a function of the initial frequency of allele B, *p*_0_, and the equilibrium frequency, *p*\*. It is demonstrated that the establishment probability largely depends on the mode of selection on allele B, which determines *p*\* (Figure 7). The establishment probability is relatively high when balancing or equalizing selection works because polymorphism at locus B/b increases the benefit of linkage. In contrast, when directional selection works (either positive or negative), linked selection does not significantly help establishment. When negative selection works, the establishment probability is low because the frequency of allele B is so low that allele A has difficulty linking with it. When positive selection works, the establishment probability is also low because beneficial allele B should be almost fixed. The effect of *p*_0_ appears to be smaller unless *p*_0_ is very different from *p*\*.

Our results demonstrate that the fate of newly arisen sex-determining mutation is mainly determined in the early phases (i.e., stochastic phase), where random genetic drift is the major evolutionary force. It is indicated that the stochastic phase plays an important role in understanding the evolutionary process of turnover of sex-determining loci. In the stochastic phase, *p*_0_ is a very important factor affecting the initial behavior of the new mutation, and the density distribution of *p*_0_ largely depends on the mode of selection at the B/b locus. The establishment probability of allele A is very low when directional selection works at the B/b locus, indicating that the presence of loci under sexually antagonistic selection could significantly enhance turnover of sex-determining loci only when sexually antagonistic selection works in the form of balancing selection.

Our theory has great implications for the rate of turnover of sex-determining loci in natural populations. An important biological question is how sexually antagonistic loci help the turnover of sex-determining loci on the genomic scale. One might think that most turnover occurs with the help of sexually antagonistic loci under balancing selection, because the establishment probability is markedly high when balancing selection works at the B/b locus (see also van Doorn and Kirkpatrick 2007, 2010). On one hand, one might consider that there may not be a large number of sexually antagonistic loci under balancing selection in a genome, so that their relative contribution might be small. Alternatively, there might be a negligible contribution of sexually antagonistic loci under other modes of selection (i.e., equalizing and directional selection). Even if the establishment probability is not high for each, on a genomic scale, their cumulative effect may not be small. To answer the question on the relative contribution of linked selection, we need to know how many sexually antagonistic loci exist in the genome, and what mode of selection is working. While empirical studies may be powerful to address this, it is also interesting to look at polymorphism data surrounding the sex-determining locus. Our simulations (Figure 8) provide insight into how to distinguish the mode of selection at a linked locus. Shortly after turnover, if divergence between male and female haplotypes is restricted in a very narrow region around the sex-determining locus, it may be likely that the sex-determining allele has become established with no help from linked selection (i.e., Myosho *et al*. (2012); Kamiya *et al*. (2012); Koyama *et al*. (2019)). On the other hand, a highly diverged region spreads surrounding the sex-determining locus, the sex-determining allele has become established together with a linked allele at a sexually antagonistic locus, or sexually antagonistic loci arose after the establishment of the sex-determining locus.

## ACKNOWLEDGMENTS

This work was partly supported by grants from the Graduate University for Advanced Studies, SOKENDAI, and Japan Society for the Promotion of Science (JSPS) to HI and TS (19H03207, 20J21760). The authors would like to thank Enago (www.enago.jp) for the English language review.

## APPENDIX

### APPENDIX A: Details of simulation model

The details of simulation model are presented. The life cycle is assumed to be in the order of mating (random genetic drift), selection, gamete production (recombination) and mutation.

First, the details of simulations for establishment probabilities are provided. Let *s*_*ijk*_ and *e*_*ijk*_ be the frequency of genotype *ijk* among sperm and egg population at the beginning of the generation, respectively, where *i* ∈ {A, a}, *j* ∈ {B, b} and *k* ∈ {X, Y}. In a mating step, *N* individuals are produced by random mating. Denote by *f*_0_(*ijk*/*lmn*) the frequency of zygotes between a sperm of genotype *ijk* and an egg of genotype *lmn. E*[*f*_0_(*ijk*/*lmn*)] is given by *s*_*ijk*_*e*_*lmn*_. According to this probability, *N* zygotes are produced by multinomial sampling. Then, sex is determined by a combination of loci A/a and X/Y (see Figure 1B and C).

During the selection step, fitness of each genotype is determined by locus B/b. Let *F* be a set of genotype that grows into a female. The fitness of genotype *X* = *ijk*/*lmn, W*(*X*), is given by

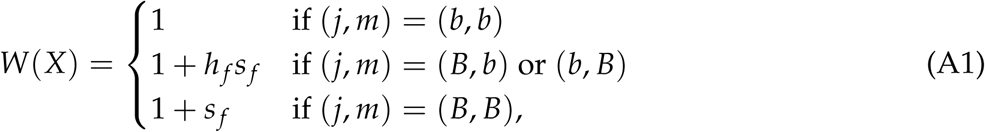

for females (i.e., *X* ∈ *F*) and

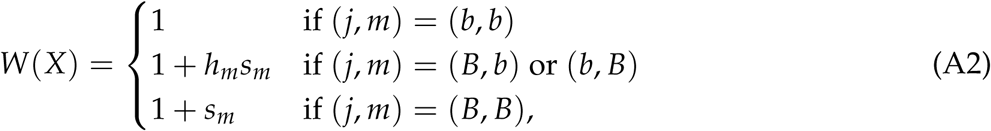

for males (i.e., *X* ∉ *F*). For *X* ∈ *F*, the genotype frequency among female population, 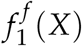, is

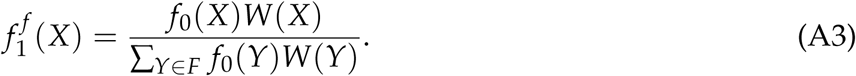

For *X* ∉ *F*, the genotype frequency among male population, 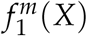, is

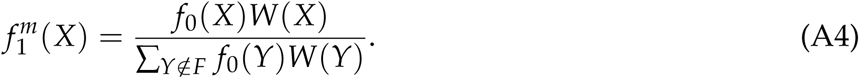

In gamete production, genotype *ijk*/*lmn* produce 8 kinds of gametes, *ijk, ijn, imk, imn, ljk, ljn, lmk* and *lmn*, with proportion 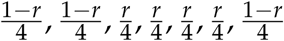 and 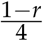, respectively. After that, proportion *u* of allele B mutates to allele b while proportion *v* of allele b mutates to allele B. Then, the frequency *s*_*ijk*_ and *e*_*ijk*_ at the beginning of the next generation is defined.

Initial condition of simulations are determined as follows. In simulations for conditional probability, initial frequency of each gamete is set as

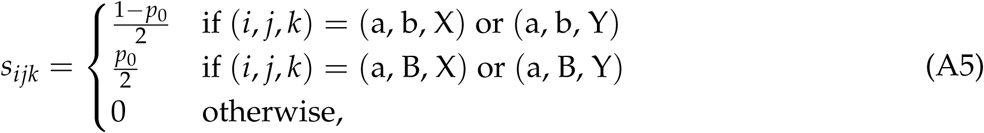

and

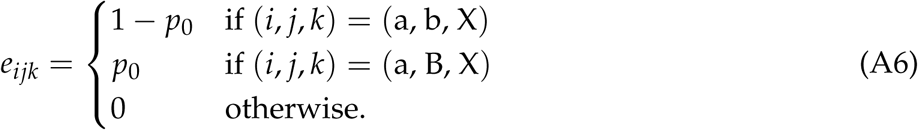

After the mating step of first generation, an allele A is introduced. To investigate the fate of allele A arising in linkage with allele *i* (*i* ∈ {B, b}), an allele *i* is randomly chosen and the linked allele a is changed into allele A. Simulation is run until either locus X/Y or locus A/a becomes monomorphic. In simulations for unconditional probability, we first run simulations without introducing allele A to obtain stationary distribution of *s*_a*jk*_ and *e*_a*jk*_. After a burn-in period of 10*N* generations, *s*_a*jk*_ and *e*_a*jk*_ are sampled. For each set of *s*_a*jk*_ and *e*_a*jk*_, an allele a is randomly chosen and turned into allele A after the mating step of first generation. Simulation is run until either locus X/Y or locus A/a becomes monomorphic.

Next, the details of simulations for the pattern of neutral polymorphism are provided. A genomic region of relative length 1 is simulated where the locus A/a is located at relative position 0.25 and the locus B/b is located at relative position 0.75. Although the basic assumptions are the same as the three loci model, implements are slightly different to improve efficiency of simulations such that all events are incorporated into the mating step.

In a mating step, a father and a mother are chosen from individuals in previous generation to form an offspring. To incorporate the selection, the probability that an individual is chosen as a parent is proportional to its relative fitness among individuals of the same sex. For each parent, a haplotype is made through recombination event where constant rate of recombination across the simulated region is assumed. By combining two haplotypes, an offspring is formed. Mutation is finally introduced in both the locus B/b and neutral sites. These procedures are repeated *N* times to generate individuals of the present generation.

The dynamics of polymorphism are simulated as follows. First, to simulate the polymorphism just before turnover, we run simulations for 200*N* generations without introducing allele A. Then, allele A is introduced at the locus A/a. If turnover occurs, simulation continues until 50*N* generations pass since the turnover.

### APPENDIX B: Derivation of Equations 1 and 15

The details for the derivation of Equations 1 and 15 are provided. First, Equation 1 is derived. We here focus on deterministic changes of allele frequency during allele A is rare and ignore stochastic changes. At the beginning of each generation, the frequencies of haplotypes A-B and A-b in the sperm population are assumed to be *x*_*B*_ and *x*_*b*_, respectively. Because locus B/b is not on the ancestral sex chromosome, we assume that the frequencies of allele B in the two sexes are almost same. By denoting the frequency of genotype *i* after random mating as *f*_0_(*i*), the frequencies of four genotypes are given by

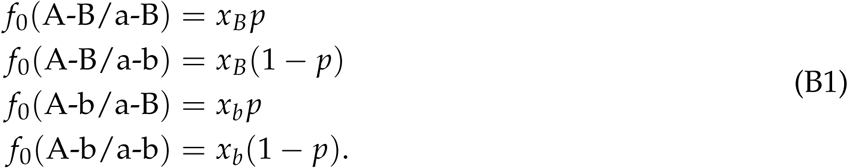

Since allele A has a masculinizing effect, the above genotypes are all male. During allele A is rare, the effect of mutants on the average fitness can be ignored. Then, the average fitness of males is 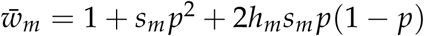. The frequency of genotype *i* after selection, *f*_1_(*i*), is given by

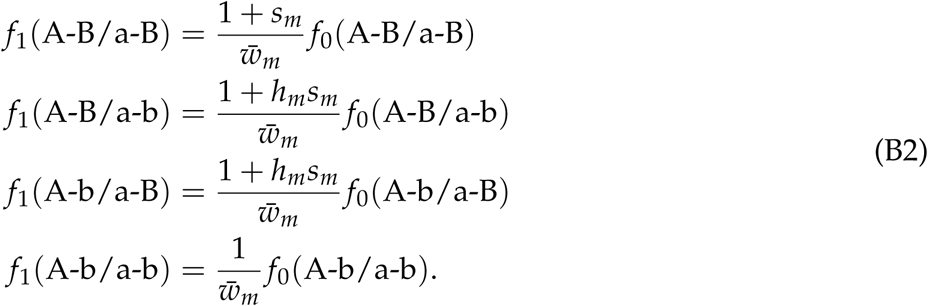

Note that a half of the population is male. Then, the haplotype frequencies after recombination among the sperm population, 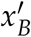 and 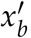, are given by

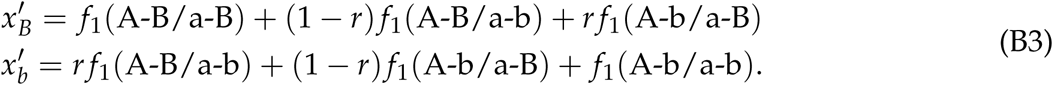

After mutation at locus B/b, the frequency changes in one generation are given by

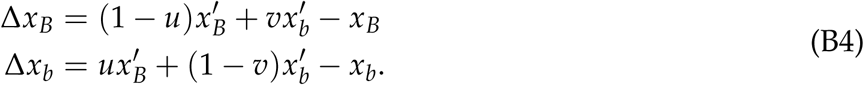

By ignoring the second order terms of *s*_*m*_, *r, u* and *v*, Equation B4 is reduced to Equation 1.

For Equation 15, the above derivations are also valid if 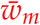, *h*_*m*_, and *s*_*m*_ are substituted by 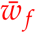, *h*_*f*_, and *s* _*f*_, because we assumed that allele A has a strong feminizing effect such that genotype XYAa grows up into females. It is obvious from the derivation that Equation 15 takes the same form as Equation 1.

### APPENDIX C: Branching process approximations

The derivation for Equations 3 and 7 are provided. Each evolutionary force is assumed to be relatively weak such that *s*_*m*_, *s* _*f*_, *r, u*, and *v* are at most the order of magnitude *ε* « 1. Denote by *λ*(*p*) the leading eigenvalue of the matrix in Equation 1. We also assume that 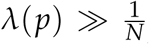 such that allele A increases deterministically once its frequency becomes large. Note that 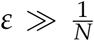 such that allele is required for 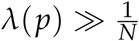.

To derive the establishment probability, we focus on descendants of an allele A after one generation. Let a random variable 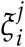 be the number of offsprings of haplotype A-*j* that is left by a haplotype A-*i* (*i, j* ∈ {B, b}). Because we assume that allele A is rare, each 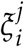 can be treated as independent and the probability distribution of 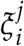 may be approximated by Poission distribution. Then, the probability generating functions of the number of offspring are given by

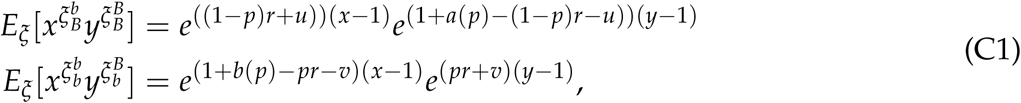

where terms of 𝒪(*ε*^2^) are ignored in the exponent. Let Δ*p* be the change of *p* in one generation. Then, establishment probabilities, *φ*_*B*_ and *φ*_*b*_, satisfy following equation:

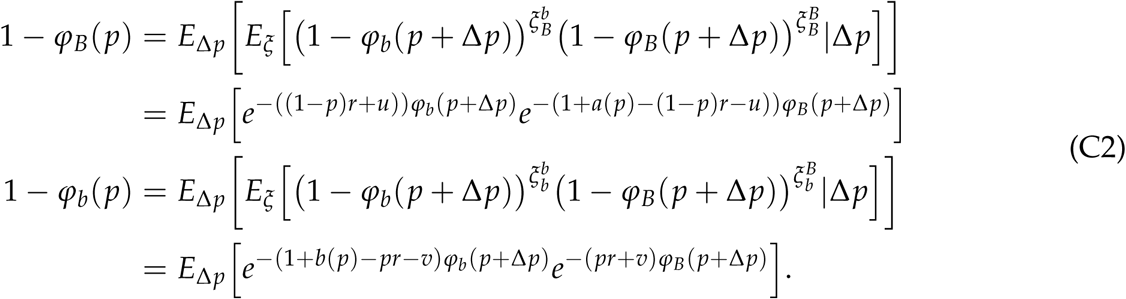

Assume that *φ*_*B*_(*p*), *φ*_*b*_(*p*) are the order of *ε*. Then, Equation C2 is expanded as

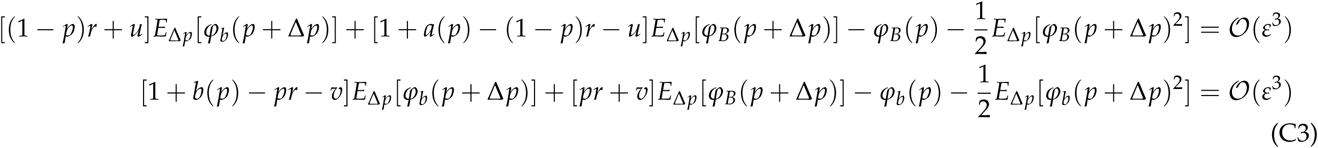

Noting that evolutionary forces are so weak that the frequency of allele B changes slowly, *E*_Δ*p*_[*φ*_*i*_(*p* + Δ*p*)] and *E*_Δ*p*_[*φ*_*i*_(*p* + Δ*p*)^2^] can be approximated by

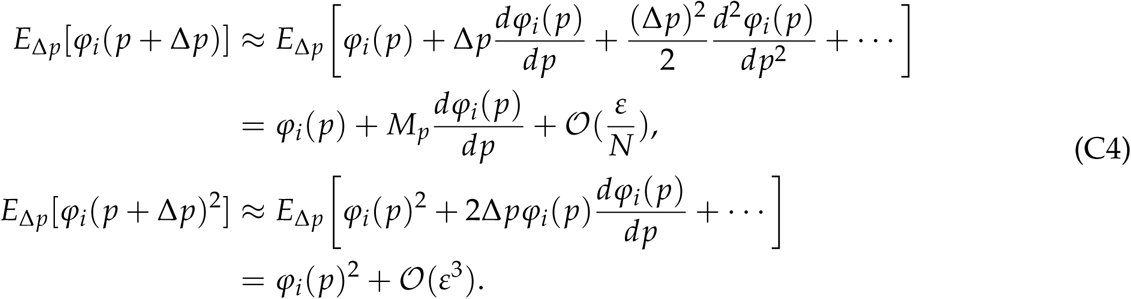

Recall that 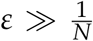 is assumed such that 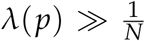 is satisfied. By substituting Equation C4 into Equation C3 and taking terms up to *ε*^2^, Equation 7 is derived. By assuming *M*_*p*_ is negligible (i.e., 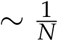), Equation 3 is also derived.

The scope and the limitation of the approximation are discussed. Equation 3 and 7 are accurate if terms of order 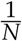 are ignored, as long as 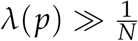. Noting that 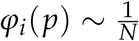 when 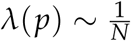, they are generally accurate if the order of 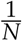 can be ignored. Since such small terms can be ignored in many cases, our approximations may be valid in broad situations.

However, there are exceptional situations, for which terms of order 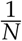 become too large to be ignored. Such situations occur when 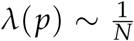 when *p* ≈ *p*\* while 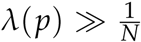 otherwise. Typical situations are that negative selection works on allele B and *r* is large, or positive selection works. In such cases, it is probable that the number of allele A may increase ∼ *N* before *p* reaches *p*\*, especially in small populations. Although *φ*_*i*_(*p*) are still proportional to 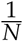, the absolute value of establishment probability may become large. For the details of these cases, see APPENDIX E.

### APPENDIX D: Explicit solution for Equation 3

The details for the derivation of Equations 4 and 5 are provided. The first equation of Equation 3 is rearranged to Equation 5. By substituting Equation 5 into the second equation of Equation 3, a quartic equation can be obtained:

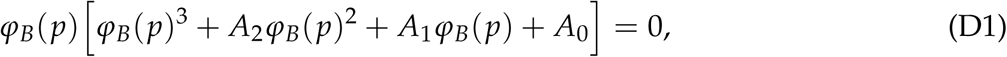

where *A*_*i*_ are defined in the main text. The largest real root of the Equation D1 is the biologically relevant solution (see also Sakamoto and Innan (2019)). The root is expressed by Equation 4 (see Abramowitz and Stegun 1970; Oldham *et al*. 2010).

### APPENDIX E: Establishment probability when linked selection does not work at *p* = *p*^*^

We describe the behavior of the establishment probability when linked selection does not work at *p* = *p*\*. This is a typical situation when negative selection works against allele B and *r* is large, or positive selection works for allele B, so that the establishment probability is very small (unless *p*_0_ is very different from *p*\*). We here explore such a case in detail because Equation 7 could underestimate the establishment probability (see Figures 5 and 6). This is because our derivation based on the branching process approximation ignores establishment events occurring in a nearly neutral fashion. In this APPENDIX, we derive another approximation focusing on the establishment of allele A through a nearly neutral fashion. It is assumed that selection is strong and *p*_0_ is close to *p*\*.

First, we consider Case 1 (i.e., the turnover without changing heterogametic sex) and negative selection on allele B is assumed. For linked selection not to work at *p* ≈ *p*\*, *r > h*_*m*_*s*_*m*_ is also assumed (see Equation 9). Under this condition, the establishment process does not occur simply by increasing the frequency of haplotype A-B: rather, because frequent recombination keeps breaking the linkage between alleles A and B, allele A has to increase without increasing allele B. In practice, negative selection works to reduce the frequency of allele B and allele b is almost fixed. Therefore, establishment occurs such that haplotype A-b increases. In this case, haplotype A-b is selectively neutral, so that we can derive an approximation of the establishment probability by following Equation 8. By using 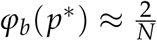, where 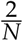 is the establishment probability of a neutral masculinizing allele, the establishment probability is approximated by

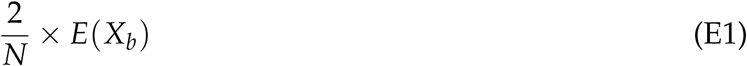

where *X*_*b*_ is the number of haplotype A-b when *p* reaches *p*\*. We can calculate *E*(*X*_*b*_) numerically under the branching process approximation as described below.

Next, positive selection on allele B is assumed in Case 1. At equilibrium with *p*\* ≈ 1, haplotype A-B is selectively neutral because the frequency of allele b is very low. Similar to the previous case, we can derive an approximate formula for the establishmemt probability as

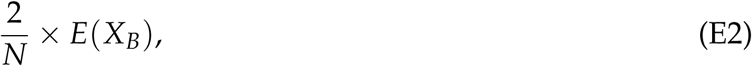

where *X*_*B*_ is the number of haplotype A-B when *p* reaches *p*\*.

We briefly explain how *E*(*X*_*B*_) and *E*(*X*_*b*_) can be computed. We here assume that mutation rate is low enough to be ignored. Let 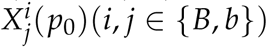 denote the number of descendant A-*j* haplotypes at *p* = *p*\* originated from the focal single haplotype A-*i* that arose when *p* = *p*_0_. *E*(*X*_*B*_) and *E*(*X*_*b*_) depend on *p*_0_ and the allele at locus B/b with which the allele A initially links. By considering the change of the haplotype frequency in one generation, we can obtain the recursion equations:

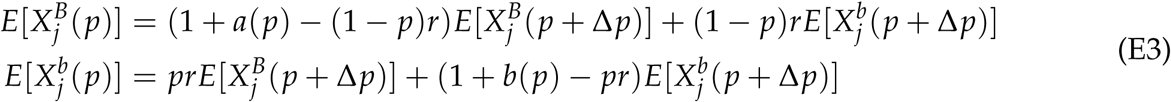

(for each coefficient, see Equation 1), where Δ*p* is the expected change of *p* in one generation. By ignoring the order of 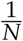 and using a continuous time approximation, Equation E3 is rearranged as

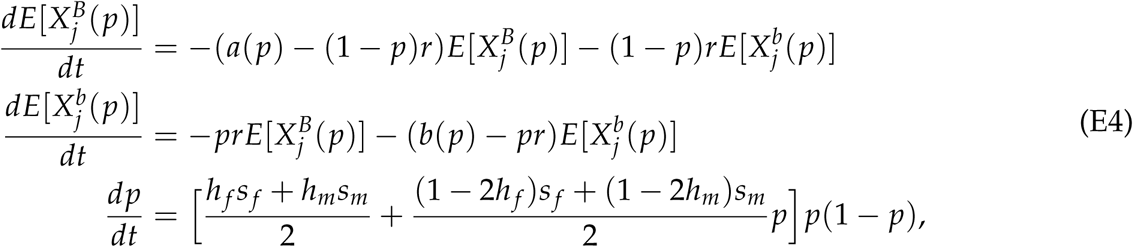

where *t* is an auxiliary variable. Then, by setting the initial condition at *t* = 0, we can obtain 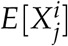 as a function of *p*.

When negative selection is assumed, the initial condition is set by the values at equilibrium *p*\* = 0 as 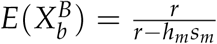 and 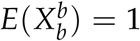. However, for a technical reason, we cannot use *p* = 0 at initial state because we cannot calculate Equation E4 under 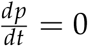 (see the main text). Instead, we set *p* = *ε* (*ε* « 1) as an initial state. When positive selection is assumed, 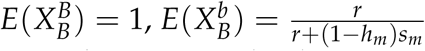 and *p*= 1 − *ε* is used as values at *t* = 0. As long as *ε* is small, its effect on the numerical value appears to be very subtle.

For the case of negative selection, the establishment probability is plotted for different population sizes *N* = 10, 000 and 100, 000 (Figure S1). All other parameters are the same as those used for Figure 5B, so that Figure S1A is identical to Figure 5B. In Figure S1, simulation results are plotted together with Equations 3, 7 and E1. Note that Equation E1 can be applied only when *r > h*_*m*_*s*_*m*_. It is demonstrated that, when *r < h*_*m*_*s*_*m*_, as stated in the main text, establishment is driven by selection such that its probability does not depend on the population size (*r* = 0.0, 0.005 in Figure S1). On the other hand, when *r > h*_*m*_*s*_*m*_, the establishment probability is no longer determined by the selection intensity, and proportional to 1/*N* because random genetic drift is the dominant force involved in the establishment. This is shown in the inner panels for *r* = 0.01, 0.02 in Figure S1, where Equation E1 agrees with the simulation results better than Equations 3 and 7 especially for a large *r* (i.e., *r* = 0.02).

Figure S2 shows the establishment probability for the case of positive selection. Simulation results are plotted together with Equations 3, 7 and E2. Two population sizes (*N* = 10, 000, 100, 000) are considered, and all other parameters are the same as those used for Figure 6 so that Figure S2A is identical to Figure 6 except that the establishment probability is log-scaled in Figure S2. For all recombination values, as the establishment process depends on random genetic drift, the establishment probability is on the order of 1/*N* (see Figure S2). Equation E2 agrees very well with simulations unless *p*_0_ is very small.

Finally, we consider Case 2 that involves a turnover of the heterogametic sex. Approximation formulae can be derived in a similar way to Case 1. By noting that the establishment probability of a neutral feminizing allele is 1.07/*N* (Veller *et al*. 2017), the establishment probability can be approximated by,

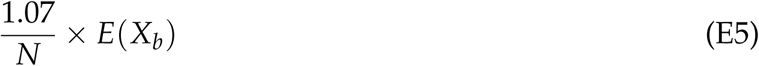

when negative selection works and *r > h* _*f*_ *s* _*f*_, and

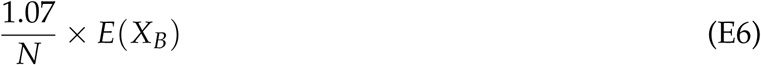

when positive selection works. *E*(*X*_*B*_) and *E*(*X*_*b*_) are calculated by using Equation E4 with substituting *h*_*m*_ and *s*_*m*_ by *h*_*f*_ and *s* _*f*_. These equations agree with simulation results (see Figure S3 and S4).

**Figure S1.**
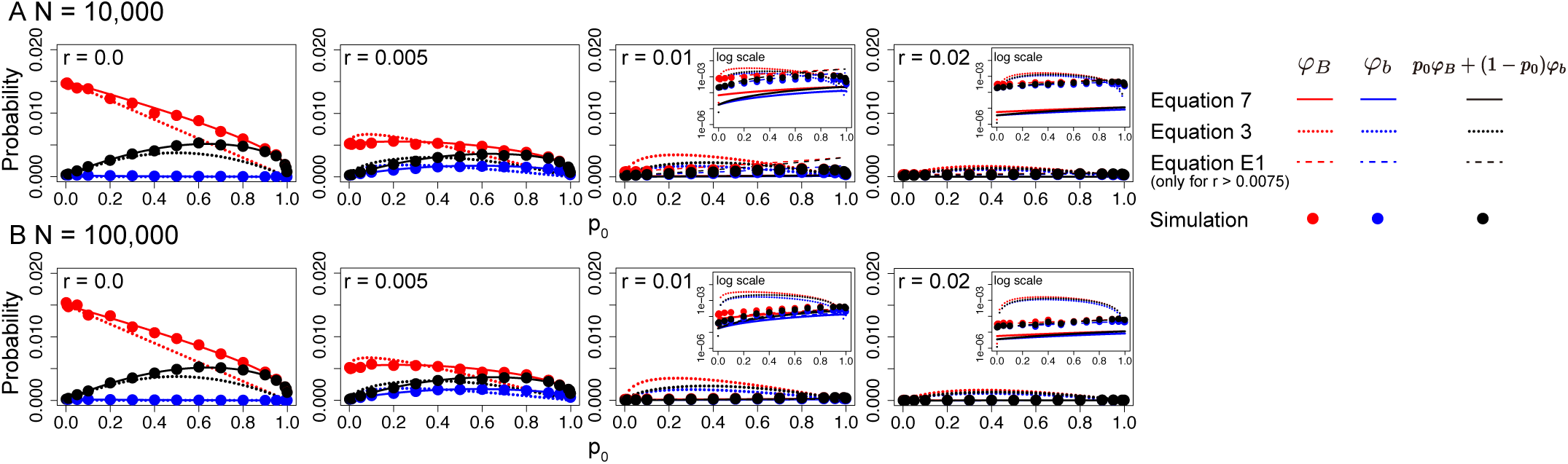
Establishment probability for the case of negative selection against allele B in Case 1. Different population sizes are assumed: (A) *N* = 10, 000, and (B) *N* = 100, 000. Other parameters are *s*_*m*_ = 0.015, *s* _*f*_ = −0.03, *h*_*m*_ = *h*_*f*_ = 0.5 and *u* = *v* = 1.0 × 10^−6^. Note that Equation E1 is plotted only for *r* = 0.01, 0.02. In the inner panels, the y-axis is log-scaled. Error bars on the red and blue circles represent the 95 % confidence interval, but they are too small to be seen.

**Figure S2.**
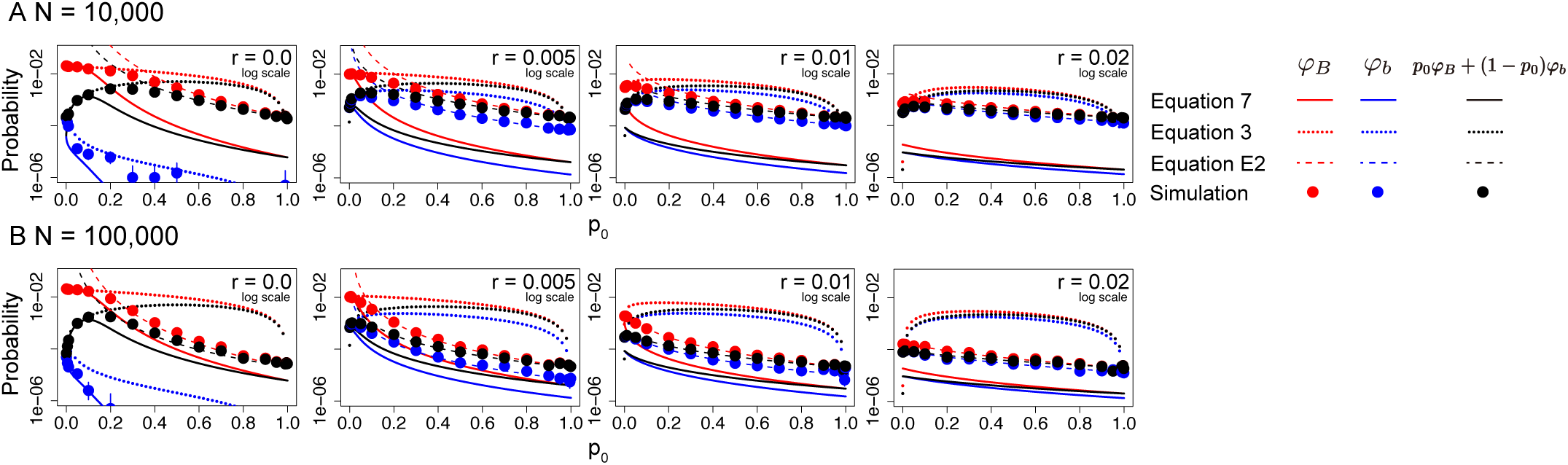
Establishment probability for the case of positive selection for allele B in Case 1. Different population sizes are assumed: (A) *N* = 10, 000, and (B) *N* = 100, 000. Other parameters are *s*_*m*_ = 0.02, *s* _*f*_ = −0.01, *h*_*m*_ = *h*_*f*_ = 0.5 and *u* = *v* = 1.0 × 10^−6^. The y-axis is log-scaled. Error bars on the red and blue circles represent the 95 % confidence interval, but they are too small to be seen.

**Figure S3.**
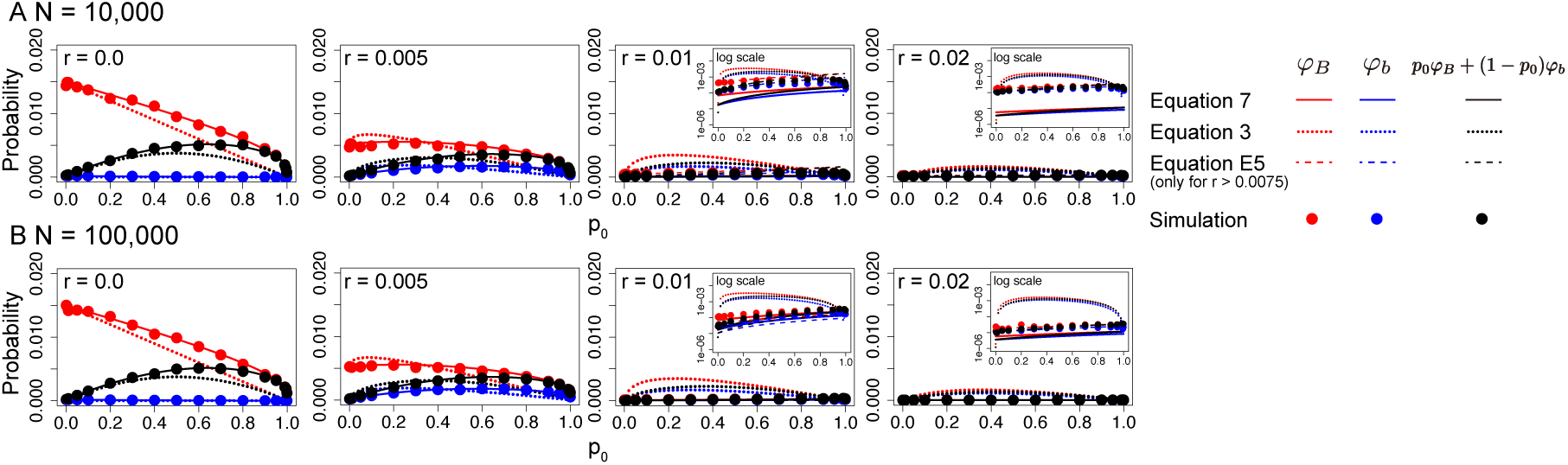
Establishment probability for the case of negative selection against allele B in Case 2. Different population sizes are assumed: (A) *N* = 10, 000, and (B) *N* = 100, 000. Other parameters are *s* _*f*_ = 0.015, *s*_*m*_ = −0.03, *h*_*m*_ = *h*_*f*_ = 0.5 and *u* = *v* = 1.0 × 10^−6^. Note that Equation E5 is plotted only for *r* = 0.01, 0.02. In the inner panels, the y-axis is log-scaled. Error bars on the red and blue circles represent the 95 % confidence interval, but they are too small to be seen.

**Figure S4.**
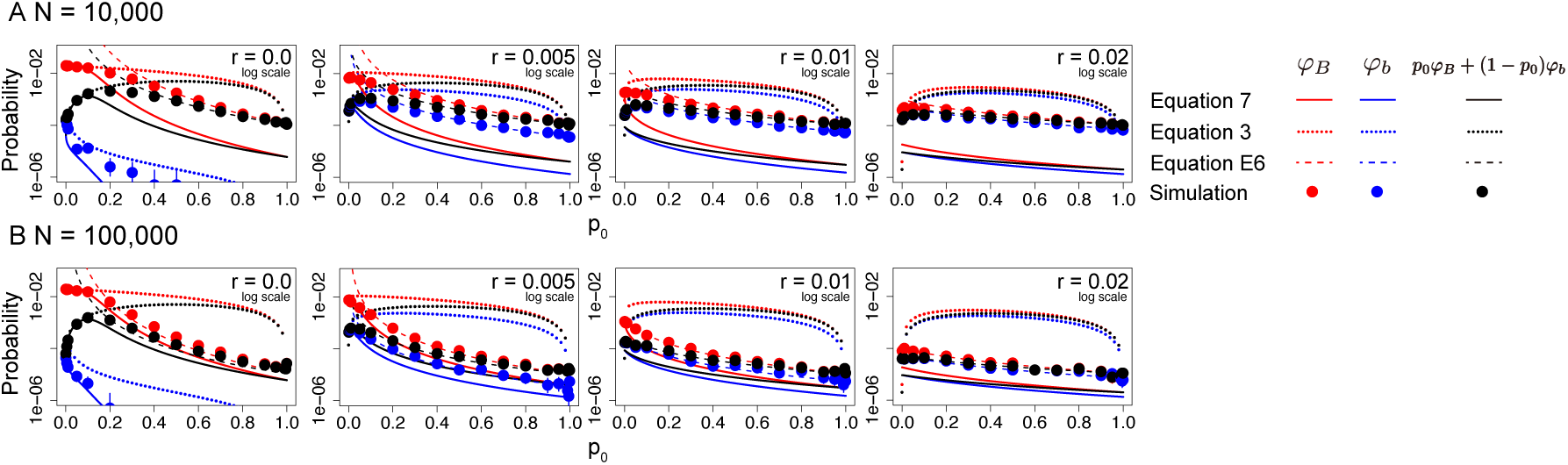
Establishment probability for the case of positive selection for allele B in Case 2. Different population sizes are assumed: (A) *N* = 10, 000, and (B) *N* = 100, 000. Other parameters are *s* _*f*_ = 0.02, *s*_*m*_ = −0.01, *h*_*m*_ = *h*_*f*_ = 0.5 and *u* = *v* = 1.0 × 10^−6^. The y-axis is log-scaled. Error bars on the red and blue circles represent the 95 % confidence interval, but they are too small to be seen.

### APPENDIX F: Relationship with van Doorn and Kirkpatrick (2007, 2010)

We here attempt to relate our results with the previous theory of van Doorn and Kirkpatrick (2007, 2010). Because their results based on a completely different model (i.e., an infinite-size population model), it is difficult to directly compare our results and theirs. To do so, we focus on our establishment probability in a finite size population, which may be comparable to the growth rate obtained by van Doorn and Kirkpatrick (2007, 2010) assuming an infinite population model. First, we briefly overview van Doorn and Kirkpatrick (2007, 2010), and then, we provide a simple, intuitive interpretation of the difference between our results and van Doorn and Kirkpatrick (2007, 2010).

#### Overview of van Doorn and Kirkpatrick (2007, 2010)

We briefly introduce the infinite-size population models of van Doorn and Kirkpatrick (2007, 2010). Initially, the X/Y locus is the sex-determining locus, at which females have genotype XX and males have genotype XY. There is another potentially sex-determining locus A/a, at which allele a is initially fixed. Then, their model considers that a new sex-determining allele A arises at locus A/a. It is also assumed that sexually antagonistic selection works at some other loci. The model allows that there can be more than one loci under sexually antagonistic selection and that the X/Y locus can also link with one of the sexually antagonistic loci, whereas our model has only one sexually antagonistic locus. In order for their results to be comparable with ours, we here set the model of van Doorn and Kirkpatrick (2007, 2010) such that there is only one sexually antagonistic locus (corresponding to the B/b locus in our model).

Under this model, van Doorn and Kirkpatrick (2007, 2010) derived the “invasion fitness” of a newly arisen allele A with considering the effect of linked selection. The invasion fitness is defined as the growth rate of allele A when the frequency of allele A is very low. They derived the invasion fitness both for a masculinizing allele (van Doorn and Kirkpatrick 2007) and for a feminizing allele (van Doorn and Kirkpatrick 2010). Although the two articles consider different situations, the formulae are in the same form if we interchange the parameters of selection in male and female, which is analogous to our results. Therefore, in the following, we arbitrarily chose to focus on the invasion of a feminizing allele, while the same argument will hold for a masculinizing allele.

Although they derived several approximations, we here focus on the tight linkage approximation because they demonstrated that it is most accurate. In Equations 5 and 8 in van Doorn and Kirkpatrick (2010), the invasion fitness of a feminizing allele was derived as

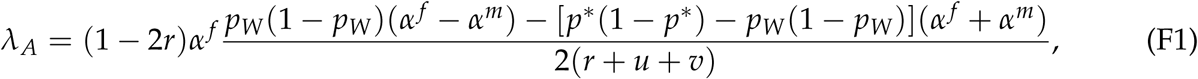

Where *α* ^*f*^ = *s* _*f*_ [*h* _*f*_ + *p* * (1 − 2*h* _*f*_)], *α* ^*m*^ = *s*_*m*_[*h*_*m*_ + *p \** (1 − 2*h*_*m*_)] and *p*_*W*_ is a solution of −(−*p \** + *p*_*W*_)*r* + *p*_*W*_ (1 − *p*_*W*_)*α* ^*f*^ + [(1 − *p*_*W*_)*v* − *p*_*W*_ *u*] = 0

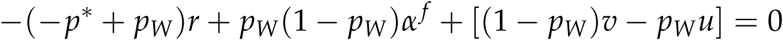

(see also the supplementary materials of van Doorn and Kirkpatrick 2010).

***Reinterpretation of van Doorn*** and ***Kirkpatrick (2007, 2010):*** We here attempt to interpret Equation F1 in the context of our stochastic model. Although the invasion fitness is based on an infinite-size population model, several studies (Connallon and Clark 2010; Yeaman and Otto 2011; Charlesworth *et al*. 2014) suggested that the establishment probability is strongly correlated with the invasion fitness, that is, the establishment probability of a single mutant allele with an invasion fitness of *λ* is given by

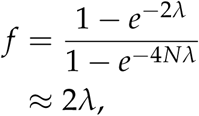

where *N* is the population size.

We find that 2*λ*_*A*_ is quantitatively in very good agreement with our *φ*_*B*_(*p*\*), the establishment probability of allele A arising in linkage with beneficial allele B, as demonstrated in Figure S5 (the blue line with + and green line with Δ in Figure S5). Note that we here use a fairly large *N* to reduce the effect of random genetic drift so that 2*λ*_*A*_ can be better understood in our framework. It appears that 2*λ*_*A*_ does not take into the case that allele A arises in linkage with allele b because 2*λ*_*A*_ does not agree with *p*\* *φ*_*B*_(*p*\*) + (1 − *p*\*)*φ*_*b*_(*p*\*) (the red line with ◯ in Figure S5). It may be concluded that their *λ*_*A*_ should well reflect the behavior of allele A when (i) the frequency of allele B is *p*\* and (ii) the focal allele A is linked with allele B, rather than the general establishment probability with no conditions that correspond to our *φ* (the black line with × in Figure S5). The first condition (i) makes sense because *p* would be at the equilibrium frequency, *p*\*, in a deterministic treatment. The second one (ii) may be understood if the case that allele A links with allele b is ignored because the invasion fitness for this case is smaller than 1.

**Figure S5.**
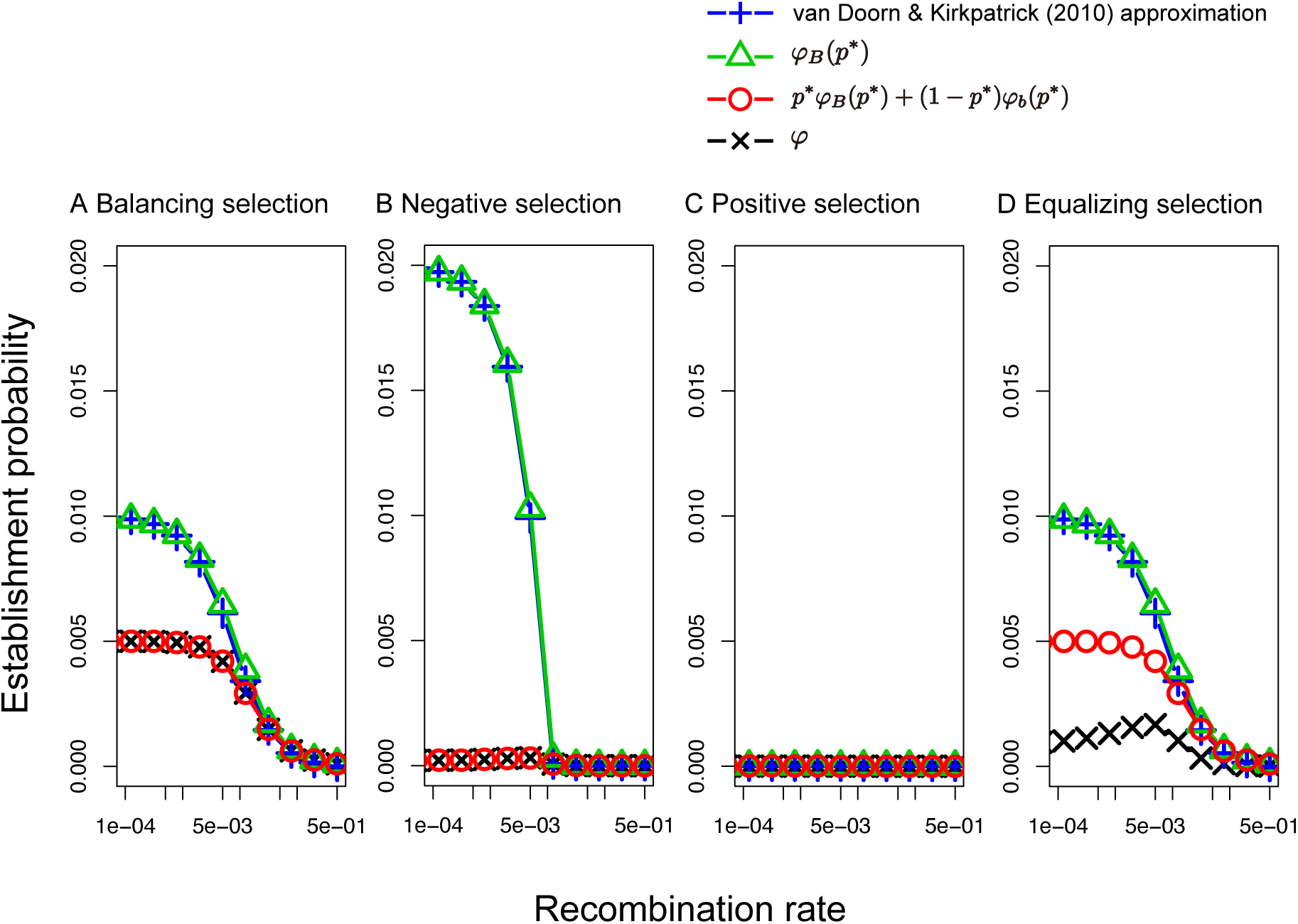
Equation 8 of van Doorn and Kirkpatrick (2010) is plotted together with our theoretical results. *N* = 100, 000 and *u* = *v* = 1.0 × 10^−6^ are assumed. Other parameters are (A) *s* _*f*_ = 0.02, *s*_*m*_ = −0.02, *h*_*f*_ = 1.0, *h*_*m*_ = 0.0, (B) *s* _*f*_ = 0.02, *s*_*m*_ = −0.025, *h*_*f*_ = *h*_*m*_ = 0.5, (C) *s* _*f*_ = 0.02, *s*_*m*_ = −0.01, *h*_*f*_ = *h*_*m*_ = 0.5 and (D) *s* _*f*_ = 0.02, *s*_*m*_ = −0.02, *h*_*f*_ = *h*_*m*_ = 0.5. Error bars on the red and blue circles represent the 95 % confidence interval, but they are too small to be seen.

**Figure S6.**
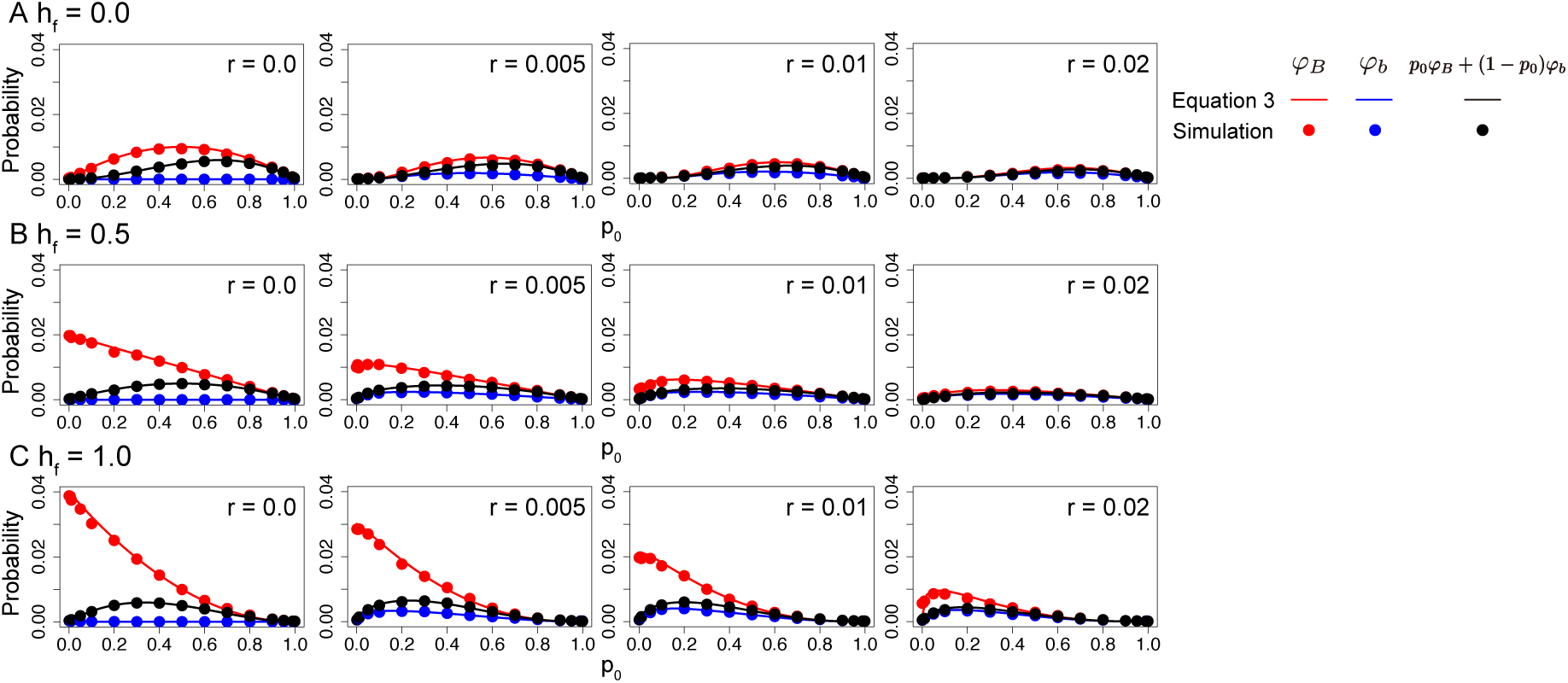
Establishment probability of allele A for different dominance and recombination rates in Case 2. Three dominance coefficients are assumed: (A) *h*_*f*_ = 0.0, (B) *h*_*f*_ = 0.5 and (C) *h*_*f*_ = 1.0. Other parameters are as follows: *s* _*f*_ = −*s*_*m*_ = 0.02, *h*_*m*_ = *h*_*f*_, *N* = 10000, *u* = *v* = 1.0 × 10^−6^. Error bars on the red and blue circles represent the 95 % confidence interval, but they are too small to be seen.

**Figure S7.**
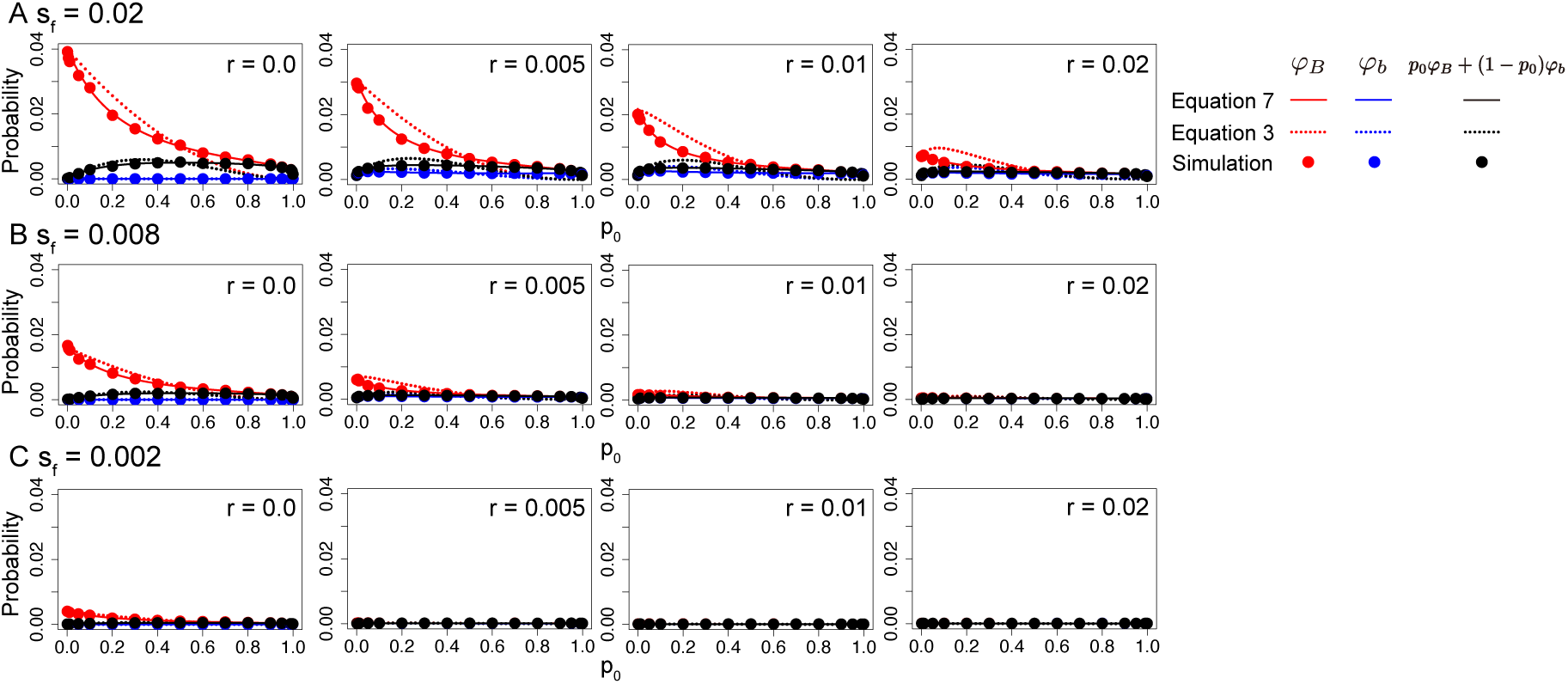
Establishment probability for the case of balancing selection for allele B in Case 2. Different strengths of selection are assumed: (A) *s* _*f*_ = −*s*_*m*_ = 0.02, (B) *s* _*f*_ = −*s*_*m*_ = 0.008 and (C) *s* _*f*_ = −*s*_*m*_ = 0.002 are assumed. Other parameters are *h*_*f*_ = 1.0, *h*_*m*_ = 0.0, *N* = 10000, *u* = *v* = 1.0 × 10^−6^. Error bars on the red and blue circles represent the 95 % confidence interval, but they are too small to be seen.

**Figure S8.**
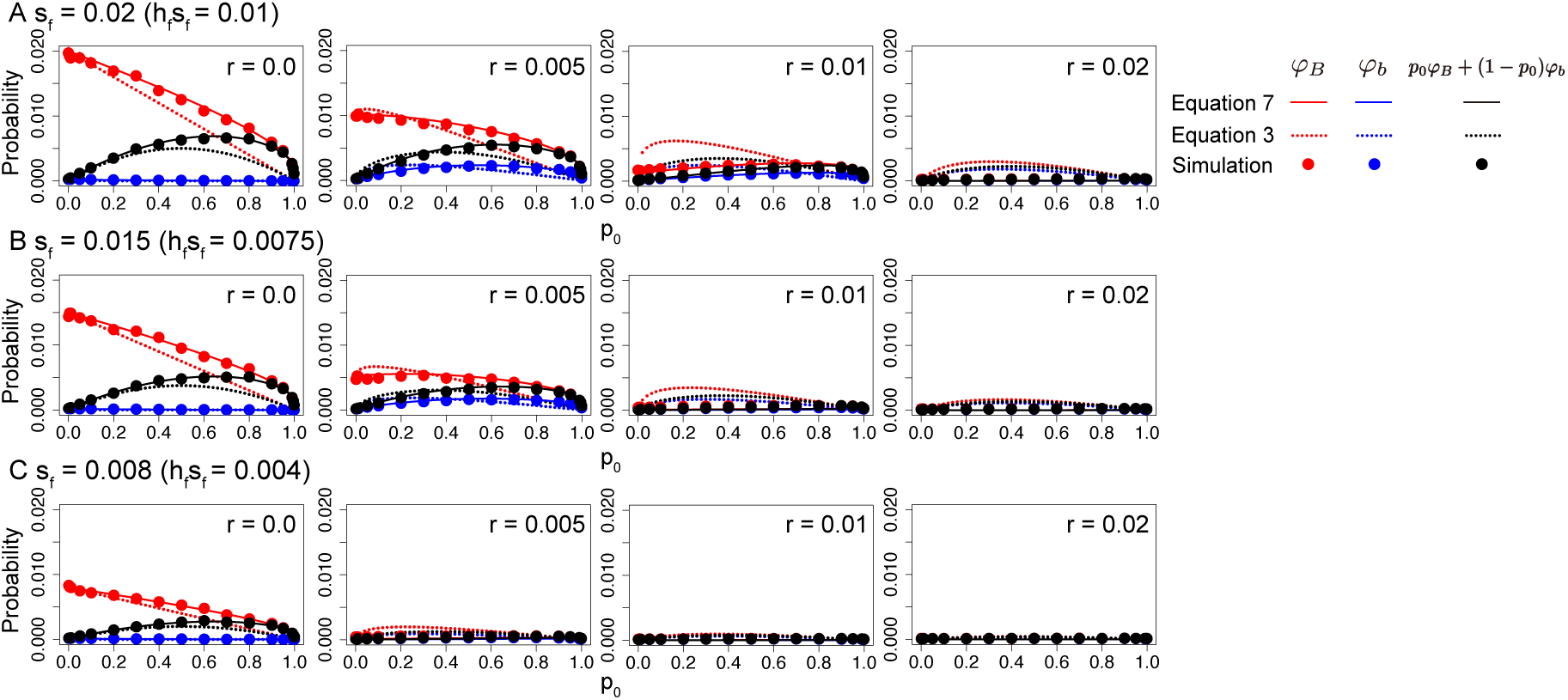
Establishment probability for the case of negative selection against allele B in Case 2. Different strengths of selection are assumed: (A) *s* _*f*_ = 0.02, (B) *s* _*f*_ = 0.015 and (C) *s* _*f*_ = 0.008. Other parameters are *s*_*m*_ = −2*s*_*f*_, *h*_*m*_ = *h*_*f*_ = 0.5, *N* = 10, 000, and *u* = *v* = 1.0 × 10^−6^. Error bars on the red and blue circles represent the 95 % confidence interval, but they are too small to be seen.

**Figure S9.**
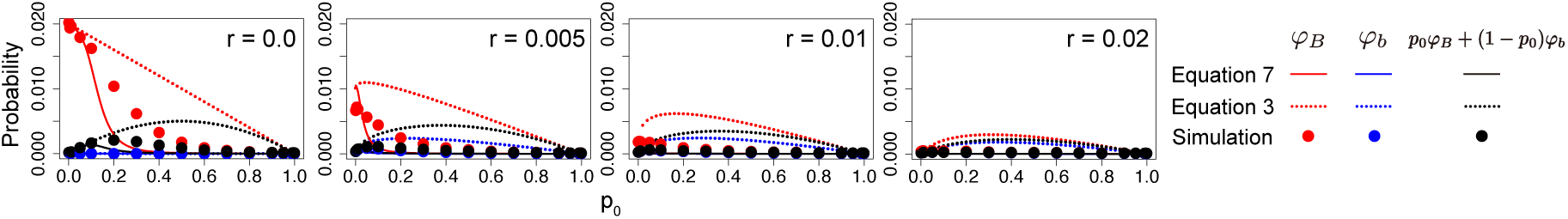
Establishment probability for the case of positive selection for allele B in Case 2. Other parameters are *s* _*f*_ = 0.02, *s*_*m*_ = −0.01, *h*_*m*_ = *h*_*f*_ = 0.5, *N* = 10, 000, and *u* = *v* = 1.0 × 10^−6^. Error bars on the red and blue circles represent the 95 % confidence interval, but they are too small to be seen.

**Figure S10.**
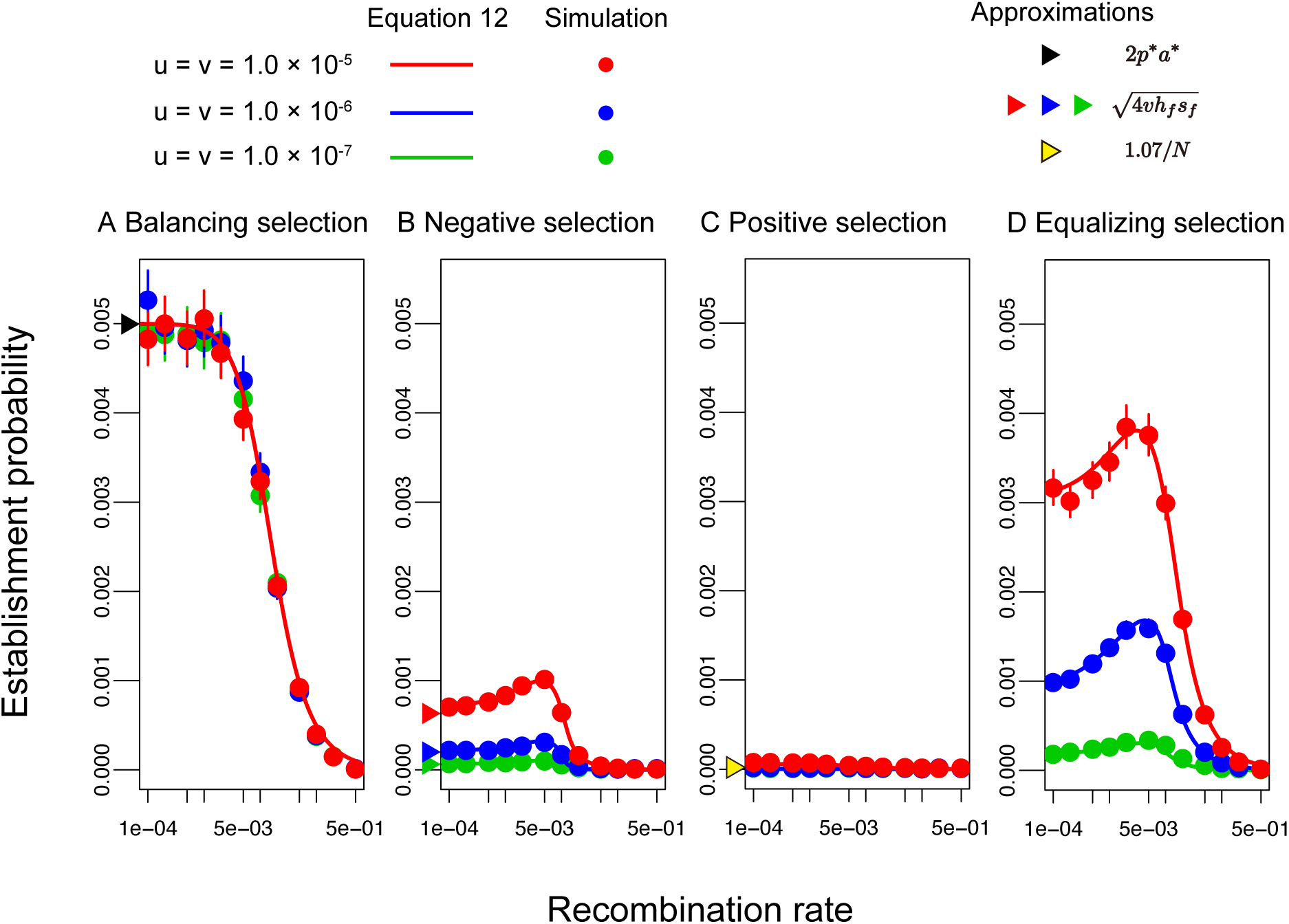
Establishment probability of a feminizing allele for different modes of sexually antagonistic selection. *N* = 100, 000 and *u* = *v* are assumed. Other parameters are (A) *s* _*f*_ = 0.02, *s*_*m*_ = −0.02, *h*_*f*_ = 1.0, *h*_*m*_ = 0.0, (B) *s* _*f*_ = 0.02, *s*_*m*_ = − 0.025, *h*_*m*_ = *h*_*f*_ = 0.5, (C) *s* _*f*_ = 0.02, *s*_*m*_ = −0.01, *h*_*m*_ = *h*_*f*_ = 0.5 and (D) *s* _*f*_ = 0.02, *s*_*m*_ = −0.02, *h*_*m*_ = *h*_*f*_ = 0.5. Error bars on the red and blue circles represent the 95 % confidence interval.

